# Calcium channel-coupled transcription factors facilitate direct nuclear signaling

**DOI:** 10.1101/2025.02.09.637126

**Authors:** Eshaan R. Rao, Tyler Thaxton, Eric Gama, Jack Godfrey, Cenfu Wei, Qiaoshan Lin, Yan Li, Daniel Parviz Hejazi Pastor, Christian Hansel, Xiaofei Du, Christopher M. Gomez

**Author notes:** These authors contributed equally to this work (Eshaan R. Rao, Tyler Thaxton).

## Abstract

VGCCs play crucial roles within the CNS, in maintaining cell excitability, enabling activity-dependent neuronal development, and forming long-term memory by regulating Ca^2+^ influx. The intracellular carboxyl-terminal domains of VGCC α1 subunits help regulate VGCC function.

Emerging evidence suggests that some VGCC C-termini have functions independent of channel gating and exist as stable proteins. Here, we demonstrate that all VGCC gene family members express bicistronic mRNA transcripts that produce functionally distinct C-terminal proteins (CTPs) in tandem with full-length VGCC α1 subunits. Two of these CTPs, α1CCT and α1ACT, cycle to and from the nucleus in a Ca^2+^- and calmodulin-dependent fashion. α1CCT, α1ACT, and α1HCT regulate chromatin accessibility and/or bind directly to genes, regulating gene networks involved in neuronal differentiation and synaptic function in a Ca^2+^-dependent manner. This study elucidates a conserved process of coordinated protein expression within the VGCC family, coupling the channel function with VGCC C-terminal transcription factors.

## Introduction

Calcium (Ca^2+^) is a ubiquitous second messenger that regulates numerous biological properties and activities throughout eukaryotic cells. Ca^2+^ signaling is especially important in the nervous system, acting both intra- and intercellularly to convey critical information necessary for its proper development and function. One tool within the neuronal Ca^2+^ signaling toolkit is the diverse family of Ca^2+^-permeable channels present among neuron subtypes, each one having unique gating and transduction pathways. These voltage-gated Ca^2+^ channels (VGCCs) are integral in maintaining neuronal excitability, synaptic transmission, muscle contractility, activity-dependent changes in gene expression, and long-term learning and memory changes within the central nervous system (CNS). Ca^2+^ signaling through VGCCs has also been shown to be critically important for proper migration of neuronal cells, such as granule cells in the cerebellum, neurons in the postnatal olfactory bulb, and interneurons in the cortex (Bortone and Polleux, 2009; Darcy and Isaacson, 2009; Komuro and Rakic, 1996).

The main pore-forming alpha 1 (α1) subunits of Ca^2+^ channels comprise a family encoded by ten distinct genes (*CACNA1X*) corresponding to three subfamilies (Ca_v_1, Ca_v_2, and Ca_v_3), and give rise to distinct high-voltage activated/high conductance (L-type, Ca_v_1), high-voltage activated/low conductance (Ca_v_2) and low-voltage activated/transient (Ca_v_3) Ca^2+^ channels. Alpha 1 subunits assemble with accessory subunits and have important interactions via their intracellular C-terminal domains with several intracellular proteins to confer gating properties, mediate signal transduction and facilitate subcellular compartmentalization (Best and Kamp, 2012; Neely and Hidalgo, 2014). Numerous loss- or gain-of-function mutations in each of the α1 subunits have wide- ranging effects on the VGCC gene products and lead to debilitating neurological, neuropsychiatric, muscle, sensory, or cardiac disorders. However, due to their phenotypic and genetic complexity, clear insights into pathogenesis and therapeutic strategies remain elusive for these genetic disorders.

Most well-recognized examples of Ca^2+^-coupled processes represent demonstrable integration between ion channels and separate signaling molecules encoded by distinct genes, such as the Ca^2+^-dependent enzymes, calmodulin-dependent protein kinase II (CaMKII), calcineurin, and protein kinase C (PKC) (Dolmetsch et al., 2001; Wheeler et al., 1994; Wheeler et al., 2008; Wheeler et al., 2012). The intracellular translocation and activation of certain transcription factors (TFs), including the developmental TFs, NFAT, CREB, and MEF-2, is regulated by highly precise spatial and temporal Ca^2+^ signaling within neurons (Beals et al., 1997; Belfield et al., 2006; Hardingham et al., 2001; Linseman et al., 2003; Mao et al., 1999; Wu et al., 2001).

Recent studies have suggested a role for Ca^2+^ channel proteins in a more direct form of nuclear signaling, through the production of Ca^2+^ channel C-terminal proteins (CTPs) (Bannister et al., 2013; Du et al., 2013; Gomez-Ospina et al., 2006; Yang et al., 2022). The origin and function of these CTPs has not been definitively ascertained, although our previous findings suggest that, for at least some cases, *e.g., CACNA1C, CACNA1A,* and *CACNA1H,* encoding α1C, α1A, and α1H respectively, this may represent a form of *genetic coupling* to neuronal Ca^2+^ signaling molecules (Du et al., 2019), as has been recognized for several other mammalian genes (Karginov et al., 2017). The *CACNA1A* mRNA, for example, encodes the α1A Ca^2+^ channel subunit of the P/Q-type VGCC Ca_v_2.1, as well as a second protein, a newly recognized TF, termed α1ACT, which activates an ensemble of genes that are critical for early cerebellar development. Mutations in *CACNA1A* have also been associated with cerebellar developmental disorders. Pathological expansion of a glutamine repeat tract in the α1ACT protein abolishes normal TF activity and mediates the progressive disease spinocerebellar ataxia type 6 (Du et al., 2013; Du et al., 2019). Furthermore, some mutations in the *CACNA1C* gene, encoding the Ca_v_1.2, α1C subunit, as well as a C-terminal TF, lead to cardiac defects, arrhythmias, and profound neurodevelopmental phenotypes (Cipriano et al., 2024; Gomez-Ospina et al., 2013; Gomez-Ospina et al., 2006; Rodan et al., 2021).

In this study, we formally explored the molecular basis, physiological control, and function of CTP expression for the VGCC gene family. Our findings demonstrate that all ten neuronal VGCC genes appear to express α1 Ca^2+^ channel CTPs by an alternative translation process, with these ten CTPs being distributed in both cytoplasm and nucleus. Detailed examination of the mRNA transcripts from the three branches of the VGCC family encoding α1C, α1A, and α1H revealed that the CTPs, referred to here as α1ACT, α1CCT and α1HCT, respectively, are generated from overlapping cistrons through a non-canonical, cap-independent mechanism. For at least two of these CTPs, α1ACT and α1CCT, nuclear translocation is functionally coupled to Ca^2+^-calmodulin signaling through ion channels in primary neurons.

Furthermore, we found that in human neural progenitor cells (hNPCs), α1CCT, α1ACT and α1HCT, each interact with DNA to regulate the expression of an array of genes crucial for proper neuronal development/differentiation and synapse formation. Interestingly, these functions are Ca^2+^-dependent and involve specific TF motif-binding in response to depolarization. Finally, when tested *in vivo*, α1CCT restores the disrupted gene expression program in a conditional *CACNA1C* knockout mouse.

These findings reveal a conserved, coordinated strategy of genetically and physiologically coupled signals for modulating gene expression among these neuronal VGCC genes. Dysregulation of this efficient strategy may explain the widespread, pleiotropic phenotypes observed in disorders related to VGCC gene mutations.

## Results

### VGCC genes encode two distinct proteins from overlapping cistrons

Based on our findings of dual protein expression by the three VGCC genes, *CACNA1C, CACNA1A,* and *CACNA1H,* we hypothesized that dual protein expression was common to the remaining members of the VGCC gene family (Du et al., 2019). To test this for the remaining seven VGCC genes, *CACNA1S* (*C*a_v_1.1), *CACNA1D* (Ca_v_1.3), *CACNA1F* (Ca_v_1.4), *CACNA1B* (Ca_v_2.2), *CACNA1E* (Ca_v_2.3), *CACNA1G* (Ca_v_3.1), and *CACNA1I* (Ca_v_3.3) (Supplementary Table 1), we transfected HEK293T cells with cDNAs expressing each of the ten VGCC α1 subunits extended at the C-terminus with a 3xFLAG epitope tag. The cDNAs corresponding to each of the VGCC genes expressed ∼250 kDa proteins detected with the anti-FLAG antibody (Figure 1D-F). In addition, these cells expressed smaller proteins that were also reactive with FLAG antibody, ranging in size from 37 - 85 kDa (Figure 1D-F; Supplementary Table 1).

**Figure 1.**
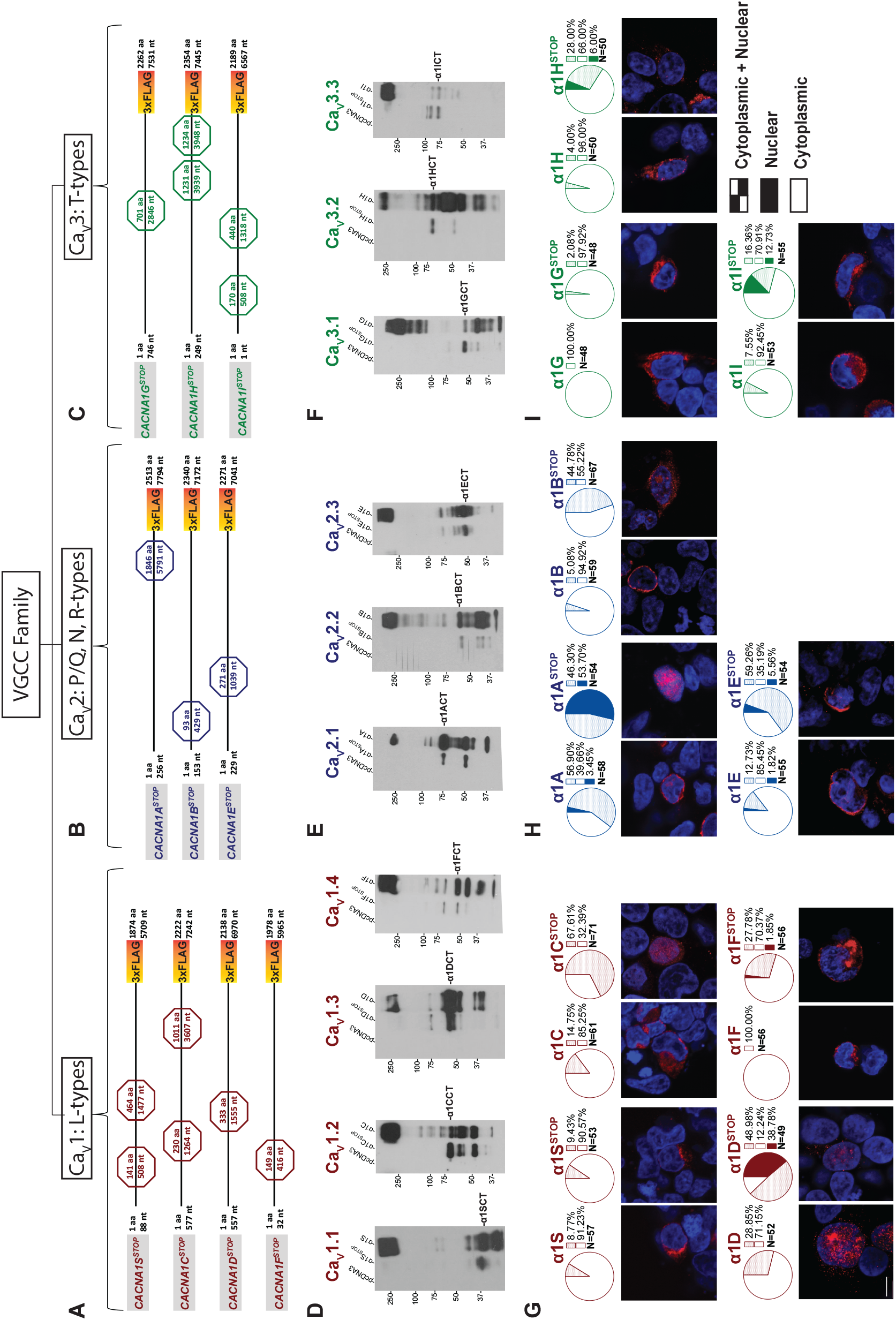
The VGCC gene family is bicistronic with C-terminal secondary proteins generated by all ten members and Subcellular localization of C-terminal secondary proteins expressed by ten VGCC cDNAs. See also Figure S1. (A) Schematic representation of L type VGCC- *CACNA1S*, *CACNA1C*, *CACNA1D*, and *CACNA1F* -with STOP codon constructs. (B) Schematic representation of P/Q, N, R type VGCC- *CACNA1A*, *CACNA1B*, and *CACNA1E* - with STOP codon constructs. (C) Schematic representation of T type VGCC- *CACNA1G*, *CACNA1H*, and *CACNA1I* -with STOP codon constructs. (D, E, and F) Western blot analysis of C-terminally 3XFLAG-tagged L type (D), P/Q, N, R type (E), and T type (F) VGCCs showing a secondary protein product detected by 3XFLAG antibody, that is produced independently of the full-length channel (FL). Secondary products ∼37-85 kDa were observed in both full length and STOP code conditions, while the primary VGCC alpha subunits (∼250 kDa) were only observed in FL conditions. Ca_V_1 sub-family (D), Ca_V_2 sub- family (E), Ca_V_3 sub-family (F). (G, H, and I) Representative confocal images of HEK293T cells transfected with either FL α1 VGCC subunit or STOP construct cDNA tagged with 3XFLAG (A, B, and C) and corresponding pie charts showing cellular compartment quantification of fluorescent signal for FL and STOP constructs in the Ca_V_1 family (G), Ca_V_2 family (H), and Ca_V_3 family (I). Number of cells used for each quantification is shown. Scale bar (G, lower left) = 10 microns.

Previous expression studies *in vitro* have attributed the presence of smaller Ca^2+^ channel fragments to proteolytic cleavage or the presence of cryptic intragenic promoters (Andrade et al., 2007; Ge et al., 2013; Gomez-Ospina et al., 2013; Gomez-Ospina et al., 2006). To demonstrate that the expression of smaller CTPs was independent of the full-length Ca^2+^ channel proteins, we inserted premature termination codons upstream of the putative CTP start sites (*CACNA1X_Stop_*), as predicted by size estimates (Figure 1A-C). In some cases, to abolish initiation at secondary methionine codons, additional stop codons were inserted. In all cases, expression of the full-length, FLAG-tagged α1 subunits was abolished and, as we had observed with *CACNA1C, CACNA1A* and *CACNA1H*, expression of the smaller 37 - 85 kDa CTPs persisted for *CACNA1S, CACNA1D, CACNA1F, CACNA1B, CACNA1E, CACNA1G,* and *CACNA1I* (Figure 1D-F). These results indicate that the CTPs are not the products of proteolytic cleavage of the full-length α1 subunits, and that expression of the CTPs in the VGCC gene family does not require expression of the full-length Ca^2+^ channel protein.

### mRNA segments of *CACNA1C, CACNA1A,* and *CACNA1H* drive internal, cap-independent translation of CTPs

We previously used protein truncation and point mutation experiments, as well as mass spectrometry, to identify the start sites for CTPs expressed by the neuronal VGCC genes, *CACNA1C*, *CACNA1A*, and *CACNA1H,* belonging to *three distinct VGCC subfamilies* and encoding α1 subunits, α1A (Ca_v_2.1), α1C (Ca_v_1.2), and α1H (Ca_v_3.2), respectively (see Figure 1A) (Du et al., 2013; Du et al., 2019). Their corresponding CTPs, α1ACT (NM_001127222.1: AUG 6114nt), α1CCT (NM_199460.3: AUG 5469nt), and α1HCT (NM_021098.2: AUG 6045nt), designated according to the α1 subunit of origin, correspond to 70 - 75 kDa proteins seen in western blots of mouse forebrain tissue probed with C-terminus-directed antisera (Du et al., 2019). To preclude the possibility that these smaller gene products arise due to the presence of cryptic promoters or alternative splicing, we used cellular transfections of *in vitro*-transcribed mRNAs. We transfected HEK293T cells with mature, polyA-tailed mRNAs encoding the three full-length VGCC α1 subunits extended at the C-terminus with a 3xFLAG epitope tag. The mRNAs were capped on the 5’ end, either with a canonical m7G cap (G-capped mRNA), allowing for initiation of translation, or with a modified 5’ cap consisting of m7G(5’)ppp(5’) A cap (A-capped mRNA). The modified A-cap inhibits binding of the initiation complex and the initiation of cap-dependent translation of the full-length α1 subunits, although may also reduce mRNA stability. In Western blots probed with an anti-FLAG antibody, we observed that G- capped mRNAs from *CACNA1C, CACNA1A,* and *CACNA1H* expressed the full-length α1 VGCC subunits of ∼250 kDa in size, as well as the smaller CTPs, α1CCT (∼70 kDa), α1ACT (∼75 kDa), and α1HCT (∼70 kDa) (Figure S1A-C). Interestingly, in cells transfected with A- capped mRNAs, the full-length α1 subunit expression was abolished while the CTPs persisted, although with reduced CTP protein expression compared to that observed with G-capped mRNAs (Figure S1A-C). We used FLAG-specific qPCR primers to confirm that the reduced CTP expression corresponded to reduced levels of A-capped VGCC mRNA, consistent with a lower stability of A-capped mRNAs, rather than an effect on translation initiation of the CTPs (Figure S1D-F). To interfere with translation of the full-length protein, we transcribed mRNAs from the cDNA templates for transfection into cells after either of two additional modifications: 1) insertion of premature termination codons between the initiation codon of the full-length α1 subunit and the putative start sites of the CTPs {*CACNA1C_stop_* (1264nt, 3607-3609nt); *CACNA1A_stop_* (5791nt); *CACNA1H_stop_* (3939 and 3941nt, 3948and 3950nt)}, or 2) insertion of sequences that generate large RNA hairpins at the 5’ end of the coding sequence, designed to impede ribosomal elongation. The RNA hairpin, with a −ΔG of −50, which is known to significantly reduce cap-dependent translation, was inserted 28 nt downstream of the T7 site in the plasmid containing the full-length VGCC open reading frame (Godfrey et al., 2022). We observed that full-length α1 subunits were not generated from mRNAs bearing either the premature termination codons or inserted hairpins, while FLAG-tagged CTPs were readily detected (Figure S1A-C), suggesting that they are generated by internal ribosomal entry, rather than a process such as ribosomal slipping or shunting.

To further verify that α1CCT and α1HCT are generated from the same mRNAs as their respective VGCC subunits, we transfected HEK293T stable cell lines expressing full-length *CACNA1C* or *CACNA1H* with siRNAs directed towards the 5’ ends of the *CACNA1C* or *CACNA1H* genes. These siRNAs abolished expression of the full-length α1 subunits as well as the secondary proteins, presumably through the RNA Induced Silencing Complex (RISC), suggesting that both are expressed from the same mRNA (Figure S1G and H). We previously reported a similar effect, induced by two distinct microRNAs, miR-711 and miR-4786-3p, directed at sequences upstream of α1ACT in cells transfected with *CACNA1A*-expressing constructs, leading to elimination of both the full length α1A and α1ACT proteins (Miyazaki et al., 2016).

Using an M-fold based algorithm of RNA folding on *CACNA1C* and *CACNA1H* VGCC gene transcripts, we predicted that the regions directly upstream of the α1CCT and α1HCT start sites form a series of highly stable stem-loop structures that could represent regions of binding by canonical translation initiation factors, as shown previously for *CACNA1A* (Du et al., 2013; Du et al., 2019). To test for the capacity of RNA segments within these regions to initiate protein translation, we inserted 1000 base pair (bp) segments of DNA from the regions directly upstream of the predicted α1CCT and α1HCT start sites into the bicistronic reporter vector pRF (*Renilla* luciferase, R-luc, and firefly luciferase, F-luc), and transfected these reporters into HEK293T cells. The coding region for R-luc is followed by a termination codon, therefore a relative increase in F-luc activity in cell lysates indicates that the inserted RNA segment can enable re-initiation of translation in a cap-independent manner (Spriggs et al., 2009). The 1000 bp segment insertions from the *CACNA1C* and *CACNA1H* genes led to significant increases in F- luc activity, 28.27±2.37 fold (p < 0.0001) and 17.38±0.14 fold (p < 0.0001) respectively, compared to empty pRF vector control (Figure S1I). These results suggest that, as with α1ACT, expression of α1CCT and α1HCT is driven by the presence of cellular IRES-like structures within the *CACNA1C* and *CACNA1H* coding regions, respectively.

Taken together, these studies indicate that the VGCC genes and their mRNAs are bicistronic, each producing the full-length α1 subunits in addition to a smaller CTP. Whereas the full-length α1 subunits are generated by canonical cap-dependent translation, the three CTPs are generated in the absence of cap-dependent translation.

### Nuclear Distribution of VGCC CTPs

α1ACT translocates to the nucleus in transfected cells (HEK293T, NIH3T3, and cerebellar granule cells), and is detected in cerebellar Purkinje cell nuclei (Du et al., 2013; Du et al., 2019; Kordasiewicz et al., 2006). To test the subcellular distribution of the other nine full-length α1 subunits and the CTP protein products, we transfected HEK293T cells with plasmids expressing each of the C-terminal 3xFLAG-tagged α1 subunits used in Figure 1, with and without inserted premature stop codons, and immunolocalized the proteins using anti-3xFLAG antibody (red fluorescence) and confocal microscopy. Immunolabeling of the cell membrane and cytoplasm was readily detectable in cells transfected with each of the intact 3xFLAG-tagged VGCC subunits (Figure 1). Interestingly, in addition to the labeling detected in the cytoplasm and membrane, nuclear staining in the same cell nucleus could be detected in cells expressing α1S, α1C, α1D, α1A, α1B, α1E, α1H, and α1I, while in α1F- and α1G-expressing cells only cytoplasmic fluorescence was detected. Since the α1 subunits are transmembrane proteins that cannot translocate to the nucleus, it is likely that proteins imaged in the nucleus are from the C- terminus. Notably, nuclear labeling was detected in cells transfected with all ten VGCC- expressing plasmids, including α1F and α1G, bearing premature stop codons (*CACNA1X_Stop_*) (Figure 1G-I). These constructs express only 3xFLAG-tagged CTPs, while expression of the full-length α1 subunit is abolished (Figure 1D-F). Also, in all cases except *CACNA1S,* the proportion of cells with labelling in the cytoplasm and membrane was decreased, while the proportion of cells with labeling in the nuclei was significantly increased when transfected with *CACNA1X_Stop_* (Supplementary Table 2). For example, for *CACNA1A*, fluorescent labelling in the nucleus was greatly increased, from an average of 63.43 ± 6.3% to 100 ± 0% (p< .05) when cells were transfected with intact *CACNA1A* compared with *CACNA1A_Stop_*, suggesting an accumulation of C-terminal 3xFLAG fusion protein. No cytoplasmic labeling was detected in cells transfected with *CACNA1A_Stop_* (Figure 1H, Supplementary Table 2). Similarly, for *CACNA1C*, nuclear fluorescence increased from an average of 13.66 ± 4.13% to 69.75 ± 16.36 % (p< .05) when cells were transfected with *CACNA1C* compared with *CACNA1C_Stop_*, again suggesting an accumulation of C-terminal 3xFLAG fusion protein. Cytoplasmic labeling reduced from an average of 86.34 ± 4.13% to 30.25% ± 16.35 % (p< .05) (Figure 1G, Supplementary Table 2). For *CACNA1H*, nuclear labeling was also greatly increased, from an average of 3.42 ± 2.15% to 37.29 ± 7.07 % (p< .05), when cells were transfected with *CACNA1H* compared with *CACNA1H_Stop_*, while cytoplasmic labeling reduced from an average of 96.57 ± 2.15% to 62.71% ± 7.07% (p< .05) (Figure 1I, Supplementary Table 2). Lastly, no nuclear labeling was detected in cells transfected with intact *CACNA1F*- and *CACNA1G*-expressing plasmids. However, for *CACNA1F_Stop_*, 62.79 ± 10.18% (p< .05) of the labeling remained in the cytoplasm, while cells with nuclear labeling increased to 37.21 ± 10.18% (Figure 1G, Supplementary Table 2). Cells transfected with *CACNA1G_Stop_* predominantly exhibited cytoplasmic labeling, with only a few cells having nuclear labeling (Figure 1I, Supplementary Table 2). In summary, increases in nuclear labeling were seen in cells expressing CTPs expressed by all ten VGCC constructs.

Together, these findings suggest that the CTPs encoded by these VGCC transcripts are present naturally in the cytoplasm and can translocate to the nucleus.

### Subcellular localization of *α*1CCT and *α*1ACT is coupled to neuronal Ca^2+^ signaling in primary cortical neurons

Given that the native CTPs are encoded by the same transcripts as the VGCC α1 subunits, and CTPs are present both in cytoplasmic and nuclear compartments, we tested whether changes in intracellular Ca^2+^ concentration, in a neuronal context, would influence CTP subcellular localization. To elicit increases in intracellular Ca^2+^, we treated primary neurons transfected with CTP-expressing constructs with either potassium chloride (KCl), glutamate (Glu), or the Ca^2+^ ionophore A23187. Exposure to increased extracellular KCl or Glu results in cellular depolarization followed by Ca^2+^ influx, partially mediated through L-type VGCCs (Sharma et al., 2008). Glu, an excitatory neurotransmitter, additionally activates glutamate-dependent cation channels (Zhang et al., 2019). A23187 is an ionophore that is highly selective for Ca^2+^ and results in rapid and robust Ca^2+^ influx in intact cells (Mattson et al., 1993; Mattson et al., 1991; Xie et al., 1996). We first studied the subcellular distribution of α1CCT, α1ACT, and α1HCT in mature primary rat cortical neurons, via fixed-cell immunocytochemistry following the induction of intracellular Ca^2+^ changes. We transfected neurons at DIV 15-17 (days in vitro) with vectors expressing 3xFLAG-tagged VGCC CTP mRNAs in culture, and manipulated intracellular Ca^2+^ concentrations with KCl (20 mM), Glu (100 µM), or A23187 (5 µM). Each treatment was evaluated in the presence and absence of the intracellular Ca^2+^ chelator, BAPTA-AM (40 µM). Neurons were treated for five minutes, after which they were fixed, and the cytoplasmic and nuclear 3xFLAG fluorescence signal for each CTP was quantified.

In untreated control neurons expressing α1CCT, the mean nuclear-to-cytoplasmic (N-C) ratio was 1.736±0.05. After treatment with 20 mM KCl, the N-C ratio was 1.955±0.07, which is slightly increased from the control (p < 0.05 vs control) (Figure 2A-C). However, exposure to Glu resulted in a significantly higher N-C ratio of 2.370±0.08 (p<0.0001 vs control) (Figure 2C). A23187 had the most pronounced effect on α1CCT nuclear-cytoplasmic localization, with an N-C ratio of 2.843±0.17 (p< 0.0001 vs control) (Figure 2C). Importantly, co-treatment of transfected neurons with the Ca^2+^ chelator BAPTA-AM during exposure to KCl, Glu or A23187 prevented the shift in N-C ratio, which was measured at 1.766±0.06 for KCl/BAPTA (p = 0.9733 vs control) and 1.861±0.09 for Glu/BAPTA (p = 0.2972 vs control), and was further reduced to 1.339±0.06 for A23187/BAPTA (p < 0.0001 vs control) (Figure 2B and C). These results indicate the importance of Ca^2+^ in the translocation of α1CCT.

**Figure 2.**
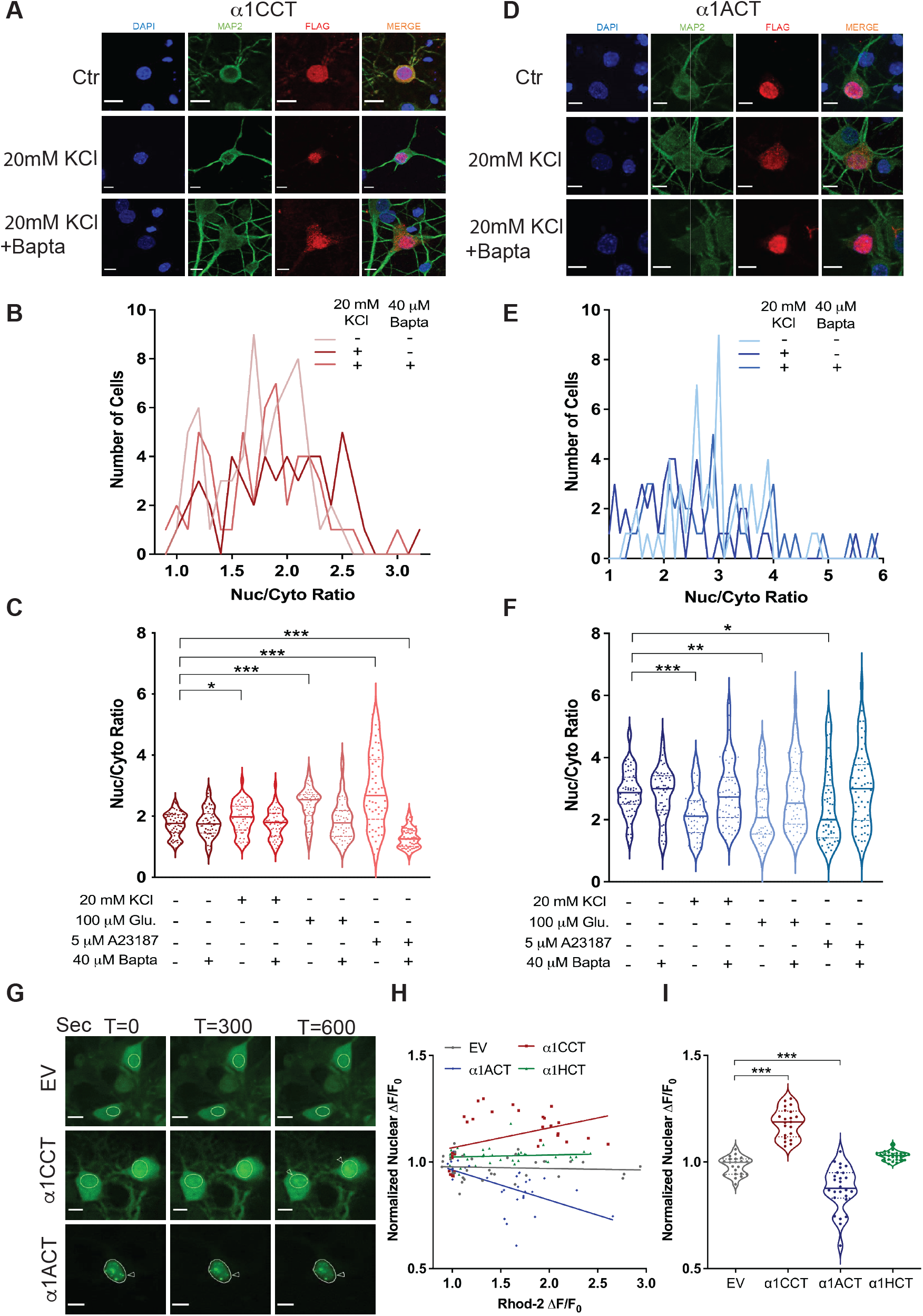
*α*1CCT and *α*1ACT translocate to or from the nucleus in response to elevated intracellular calcium. See also Figure S2. (A) Representative images of fixed rat cortical neurons transfected with α1CCT mRNA and treated with 20mM KCl with or without BAPTA-AM. Fixed neurons were stained with DAPI (blue), MAP2 antibody(green), and 3XFLAG antibody (red). Scale bars = 10 microns. (B) Quantification of nuclear/cytosolic fluorescence signal of rat cortical neurons transfected with α1CCT mRNA with 20mM KCl with or without BAPTA-AM. (C) Quantification of nuclear/cytosolic fluorescence signal of rat cortical neurons transfected with α1CCT mRNA with different treatments. (D) Representative images of fixed rat cortical neurons transfected with α1ACT mRNA and treated with 20mM KCl with or without BAPTA-AM. Fixed neurons were stained with DAPI (blue), MAP2 antibody (green), and 3XFLAG antibody (red). Scale bars = 10 microns. (E) Quantification of nuclear/cytoplasmic fluorescence signal of rat cortical neurons transfected with α1ACT mRNA with 20mM KCl with or without BAPTA-AM. (F) Quantification of nuclear/cytosolic fluorescence signal of rat cortical neurons transfected with α1ACT mRNA with different treatments. (G) Representative live-cell images of rat cortical neurons infected with AAV9-EmGFP, AAV9- α1CCT-EmGFP, or AAV9- α1ACT-EmGFP virus at T = 0 seconds, T = 300 seconds, or T = 600 seconds post-glutamate uncaging. Circles denote the nucleus. Arrows denote putative nuclear speckle formation (α1CCT) or dissipation (α1ACT). Scale bars = 10 microns. Glutamate uncaging occurred at T = 15s with two 10ms pulses of 405 nm light. Images were collected every 3s for 10 minutes total. (H) Graph depicting intracellular calcium spike (as measured by Rhod2 ΔF/F_0_) versus neuronal nuclear fluorescence change at T = 600s (ΔF/F_0_) for EmGFP, α1CCT, α1ACT, and α1HCT infected neurons (N > 30 cells for each condition) (I) Normalized nuclear fluorescence ΔF/F_0_ for EmGFP, α1CCT, α1ACT, and α1HCT infected neurons at T=600s post-glutamate uncaging (N > 30 cells per condition, *p<0.05, **p<0.01, ***p<0.001).

For α1ACT, activity-mimicking stimuli had the opposite effect on subcellular localization. In untreated control neurons expressing α1ACT, the N-C ratio was 2.88±0.10. After treatment with 20 mM KCl, the N-C ratio was reduced to 2.33±0.14 (p < 0.001 vs control) (Figure 2D-F). Treatment with Glu led to a similar reduction in the N-C ratio to 2.32±0.13 (p < 0.01 vs control), and A23187 treatment led an N-C ratio of 2.42±0.17 (p < 0.05 vs control) (Figure 2F). Moreover, the activity-induced reduction in the N-C ratio of α1ACT was prevented or reversed by Ca^2+^chelation with BAPTA-AM (Figure 2E and F), again pointing to the importance of Ca^2+^ in the translocation of α1ACT. Interestingly, none of these brief stimuli had a significant effect on the subcellular localization of α1HCT (Figure S2A - C), suggesting that the localization and translocation of the *CACNA1H*-derived CTP may not be regulated by Ca^2+^ signaling under the conditions or time course of this study. In summary, these observations demonstrate that the subcellular translocation of α1CCT and α1ACT shifts in response to activity-mimicking stimuli, suggesting Ca^2+^-dependence.

### Real-time nuclear translocation of *α*1CCT and *α*1ACT parallels Ca^2+^ signaling in primary cortical neurons

To visualize the effects of elevated intracellular Ca^2+^ on nuclear-cytoplasmic translocation of α1CCT and α1ACT in real-time, we replaced the C-terminal 3xFLAG tag with Emerald GFP (EmGFP) and monitored CTP localization and translocation in response to stimuli using live-cell confocal microscopy. Simultaneously, we used the high-affinity Ca^2+^ indicator, Rhod-2 AM, to follow intracellular Ca^2+^ levels. Primary neurons were infected with Adeno-associated virus (AAV9) expressing either α1CCT-EmGFP, α1ACT-EmGFP, or α1HCT-EmGFP, or with control empty vector expressing EmGFP (EV-EmGFP) (all driven by the EF1α promoter). Five days post-infection, neurons were loaded with Rhod-2 AM and then treated with 4-methoxy-7- nitroindolinyl (MNI) L-glutamate, a form of caged glutamate (Glu). Neuronal nuclei were identified by the fluorescent nuclear marker DRAQ5. Uncaging of Glu extracellularly around the basal dendrites and cell bodies of cortical pyramidal neurons resulted in reproducible, high-quality Ca^2+^ spikes within the targeted subset of neurons, as judged by Rhod-2 AM (red fluorescence), and these Ca^2+^ spikes frequently propagated to the nucleus (Figure S2D-F). For cells exhibiting Ca^2+^ spikes, the translocation of CTPs between nuclear and cytoplasmic compartments was recorded over a 10-minute observation period after Glu uncaging. The extent of CTP translocation into the nucleus was compared with the behavior of EV-EmGFP (and untreated CTPs-EmGFP). For α1CCT-expressing cells, Glu uncaging resulted in an *increase* in nuclear fluorescence of 20.95±1.5% (p<0.0001 vs untreated) over 10 minutes (Figure 2G-I, Supplementary Video 1 and 2). In contrast, for α1ACT-expressing cells, Glu stimulation over the same period led to a *decrease* in nuclear fluorescence of 10.73±1.6% (p<0.0001 vs untreated) (Figure 2G-I, Supplementary Video 1 and 3). Finally, while α1HCT showed no significant movement over this interval, it tended to enter the nucleus at a slower rate following intracellular Ca^2+^ spikes (Figure 2H and I, Supplementary Video 1 and 3). These results are consistent with our findings in fixed primary neurons using Glu stimuli and suggest that the translocation of α1CCT and α1ACT is dynamically coupled to Ca^2+^ levels.

### Active nuclear translocation of *α*1CCT and *α*1ACT requires Ca^2+^ signaling through VGCCs and NMDA receptors

To explore the sources of Ca^2+^ involved in mediating nuclear translocation of the CTPs, we compared the effects of distinct ion channel and receptor antagonists (Supplementary Table 3). To determine whether Ca^2+^ entry through α1C/Ca_v_1.2, α1A/Ca_v_2.1, or α1H/Ca_v_3.2 was necessary for the subsequent translocation of α1CCT or α1ACT (Adams et al., 1990; Follesa and Ticku, 1996; Hagenston et al., 2009; Kraus et al., 2010; Ruiz et al., 2009; Weiss et al., 1990), we loaded primary cortical neurons, infected with AAV9 expressing α1CCT-EmGFP or α1ACT- EmGFP, with blockers specific for each VGCC subtype, and then stimulated these neurons by Glu uncaging. Compared to Glu uncaging alone, α1CCT nuclear translocation was decreased by Ca_v_2.1 blockade (500 nM ω-agatoxin; -7.14 ± 0.021%, p = 0.0012) and by Ca_v_3.2 blockade (100 µM TTA-A2; -5.53 ± 0.020%, p = 0.0071) (Supplementary Table 4). However, the Ca_v_1.2 blocker, nifedipine (10 μM), had the most prominent effect, reducing α1CCT nuclear entry when compared to Glu uncaging alone (-15.24 ± 0.014%, p < 0.0001), suggesting that Ca^2+^ signaling through L-type VGCCs plays a prominent role in α1CCT nuclear translocation (Figure 3B-C).

**Figure 3.**
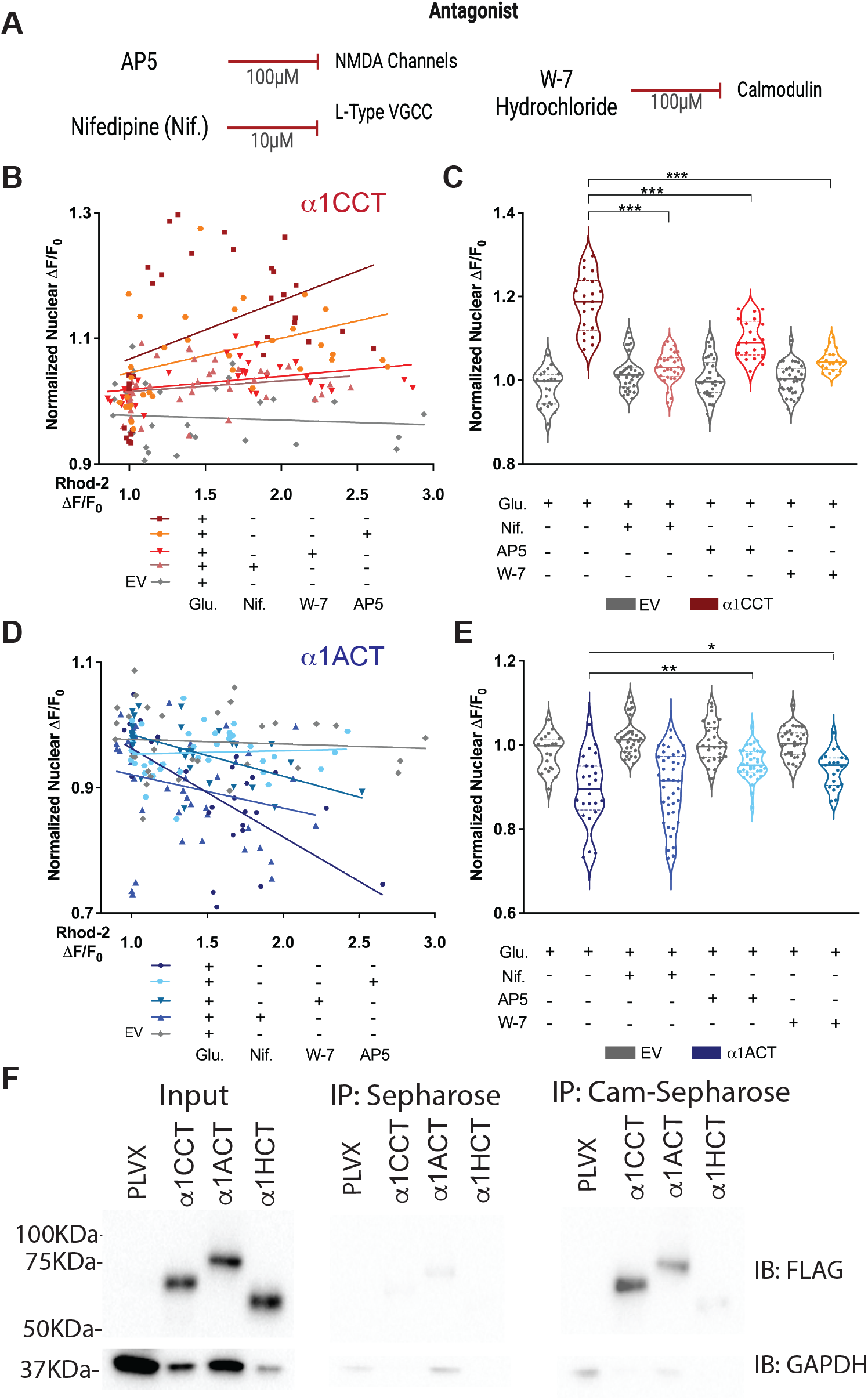
Nuclear translocation of *α*1CCT and *α*1ACT is coupled to calcium signaling through L-type VGCCs or NMDA receptors and calmodulin. See also Figure S3. (A) The calcium-channel antagonists used in conjunction with live-neuron glutamate uncaging. (B) Graph showing the comparison of α1CCT nuclear translocation following glutamate uncaging in the presence of L-type calcium channel antagonist (nifedipine), NMDA receptor antagonist (AP5), or calmodulin antagonist (W-7 hydrochloride). Negative control (-Glu) refer to Figure S2. (C) Quantification of normalized nuclear ΔF/F_0_ at T = 600 seconds for α1CCT treatment conditions. (D) Graph showing the comparison of α1ACT nuclear translocation following glutamate uncaging in the presence of nifedipine, AP5, or W-7. Negative control (-Glu) refer to Figure S2. (E) Quantification of normalized nuclear ΔF/F_0_ at T = 600 seconds for α1ACT treatment conditions. (F) Input samples showing 3XFLAG-tagged α1CCT, α1ACT, and α1HCT expression in lysates from H293T cell transfected with pLVX (control), α1CCT, α1ACT, and α1HCT. Immunoblotting (IB) was performed with anti-3XFLAG antibody, and anti-GAPDH antibody as loading control. Negative control immunoprecipitation (IP) using unconjugated Sepharose beads, showing minimal non-specific binding of α1CCT, α1ACT, and α1HCT. Calmodulin-Sepharose IP demonstrating specific interaction of α1CCT and α1ACT with calmodulin. 3XFLAG-tagged proteins were detected in the pull-down fractions, confirming their binding to calmodulin. N > 30 cells per condition, *p<0.05, **p<0.01, ***p<0.001.

To determine whether Ca^2+^ entry through glutamate-binding N-methyl-D-aspartate (NMDA) receptors contributed to α1CCT translocation, we applied the NMDA receptor antagonist, AP5 (100 µM). α1CCT nuclear entry with NMDA receptor blockade was reduced when compared to Glu uncaging alone (-6.95±0.014%, p < 0.0001), although to a lesser extent than was observed for Ca_v_1.2 blockade via nifedipine (Figure 3B-C). These results indicate that Ca^2+^ influx through either L-type VGCCs or NMDA channels may facilitate α1CCT nuclear translocation.

In neurons infected with AAV9 expressing α1ACT-EmGFP, translocation of α1ACT out of the nucleus was *unperturbed* by the presence of a VGCC blocker: 10 µM nifedipine (+0.7 ± 0.020%, p = 0.7211), 500 nM ω-agatoxin (1.94 ± 0.021%, p = 0.3551), or 100 µM TTA-A2 (1.41 ± 0.020%, p = 0.4910) (Supplementary Table 3). To identify the source of the Ca^2+^ responsible for α1ACT nuclear export, we used several additional antagonists in tandem with Glu uncaging. To investigate the role of store-operated Ca^2+^, we loaded α1ACT-EmGFP-expressing neurons with the IP3 antagonist 2-APB (50 µM) and ryanodine (100 µM), prior to Glu uncaging. α1ACT nuclear export was unperturbed following antagonism of intracellular Ca^2+^ store release via 2-APB and ryanodine (2.76 ± 0.018%, p = 0.1293 vs Glu alone) (Supplementary Table 3). However, when we exposed α1ACT-EmGFP-expressing neurons to AP5, Glu-induced α1ACT-EmGFP translocation out of the nucleus was reduced to - 4.67 ± 0.038%, compared with 10.73 ± 1.6% in the absence of AP5 (p < 0.01) (Figure 3D-E). These results suggest that Ca^2+^ entry through NMDA receptors plays a prominent role in triggering translocation of α1ACT from the nucleus.

Taken together, our results indicate that α1ACT and α1CCT, the CTPs encoded by overlapping cistrons in the mRNAs expressing α1A and α1C, respectively, are sensitive to cytoplasmic Ca^2+^ levels, and undergo dynamic nuclear-cytoplasmic shuttling within neurons in response to physiological shifts in Ca^2+^ resulting from influx through different Ca^2+^-permeable channels.

### Calcium-dependent nuclear translocation of *α*1CCT and *α*1ACT acts through calmodulin

The Ca^2+^-binding protein, calmodulin, serves as the primary intracellular receptor for Ca^2+^ and facilitates Ca^2+^-dependent signaling mechanisms in neurons (Chemin et al., 2017; Deisseroth et al., 1998; Dolmetsch et al., 2001). To test whether calmodulin affects CTP nuclear transport, we used the cell-permeant calmodulin inhibitor, W-7 hydrochloride (W-7), in primary cortical neurons transfected with GFP-tagged CTPs. W-7 reduced activity-induced α1CCT nuclear entry to 7.25±0.5%, compared with 18.34±1.5% for Glu treatment only (p<0.0001) (Figure 3B-C).

Similarly, W-7 reduced activity-induced α1ACT nuclear export to 5.76±0.9%, compared with - 10.73±1.6% for Glu treatment only (p = 0.0412) (Figure 3D-E).

Given the effect of W-7 on Ca²⁺-mediated nuclear and cytoplasmic translocation of α1ACT and α1CCT, we investigated whether there is a direct interaction between calmodulin and the CTPs. We exposed lysates from HEK293T cells expressing 3xFLAG-tagged α1CCT, α1ACT, or α1HCT, to Sepharose beads, either unconjugated or conjugated to calmodulin, and subjected precipitates to SDS-PAGE and immunoblotting. Figure 3F shows that Sepharose-calmodulin was able to precipitate 3xFLAG-tagged α1CCT (70 kDa) and α1ACT (75 kDa), although no signal was seen with lysates from cells expressing α1HCT or with Sepharose beads alone. These results suggest that the Ca^2+^-dependent nuclear-cytoplasmic translocation of α1CCT and α1ACT requires the participation of calmodulin.

### VGCC CTPs have restricted mobility within the nucleus of cultured primary neurons

Although we observed a fraction of the CTPs to cycle across the nuclear membrane, we wondered whether they also exhibited restricted mobility within the nucleus, through potential interactions with DNA, as described with some TFs (Goldstein and Hager, 2018; Govindaraj et al., 2019). To test this, we performed fluorescence recovery after photobleaching (FRAP) (Leake, 2018; Stenoien et al., 2001). We infected cultured primary neurons at DIV10 with AAV9 virus expressing EV-EmGFP, α1CCT-EmGFP, α1ACT-EmGFP, or α1HCT-EmGFP. Images were taken at DIV 15-17 following a brief (100 ms), high-intensity pulse of light to bleach the nucleus of imaged cells. We measured the recovery of fluorescence signal for 10 minutes post-bleach. Compared with EV-EmGFP alone, α1CCT-EmGFP, α1ACT-EmGFP, and α1HCT- EmGFP all exhibited significantly slower recovery times (EV t_1/2_ = 185±24s, α1CCT t_1/2_ = 532±50s, α1ACT t_1/2_ = 557±55s, α1HCT t_1/2_ = 996±55s) (Figure S3A). Additionally, quantification of the mobile fraction (MF) shows significantly lower MFs for α1CCT-EmGFP (46.7±15.9%, p < 0.01 vs. EmGFP), α1ACT-EmGFP (28.5±1.0%, p < 0.0001 vs. EmGFP), and α1HCT-EmGFP (32.5±5.3%, p < 0.0001 vs. EmGFP), compared to EmGFP (58.0±2.5%) (Figure S3B). These findings demonstrate that α1CCT-EmGFP, α1ACT-EmGFP, and α1HCT-EmGFP have a reduced nuclear mobile pool available for active nuclear transport compared to EmGFP, suggesting potential interactions with chromatin by all three CTPs.

### VGCC CTPs bind to genomic DNA in hNPCs in a calcium-dependent manner

We looked for further evidence of direct interactions of CTPs with chromatin using the CUT&RUN-seq technique (<u>C</u>leavage <u>U</u>nder <u>T</u>argets <u>&</u> <u>R</u>elease <u>U</u>sing <u>N</u>uclease, followed by high throughput sequencing). We performed the genomic binding studies in hNPC stable lines expressing either α1CCT, α1ACT, or α1HCT. The three 3xFLAG-tagged CTPs and their DNA complexes were each isolated from these hNPC stable lines, following binding by an antibody against 3xFLAG, H3K4me3 or IgG, and excision by in protein A-protein G-micrococcal nuclease (pAG-MNase). These experiments were performed under basal and depolarizing conditions, the latter induced by 20 mM KCl to elevate intracellular Ca²⁺ levels. Intracellular Ca^2^⁺ changes were confirmed using the Ca²⁺ indicator Rhod-2 AM, which showed a sustained 1.81± 0.67-fold change (p<0.003) in control hNPCs (Figure S4A) and stable nuclear CTP signals after 20 minutes of incubation. The genomic binding profiles revealed that, in either depolarized or resting conditions, α1CCT, α1ACT, and α1HCT were highly enriched within ±3000 bp of annotated transcription start sites (TSSs) (Figure 4A). In particular, α1ACT and α1HCT exhibited enhanced binding near TSS in depolarized cells compared to resting cells (Figure 4A). Binding at TSS was prominently detected in H3K4me3 antibody-isolated samples across all three CTP-expressing hNPC lines (Figure S4B), with 207 binding sites overlapping among the three CTPs (Figure S4C). In contrast, no more than one overlap in binding sites was observed among the CTP-specific samples isolated by the 3xFLAG antibody across three CTPs in either condition (Figure 4C). Comparing binding sites under depolarized and resting conditions in all three CTP expressing hNPCs, a more than 75% overlap was observed in H3K4me3 antibody-isolated samples (Figure S4D). However, the overlap in binding sites was significantly lower for 3xFLAG antibody isolated samples: α1CCT (5.93%), α1ACT (19.9%), and α1HCT (6.1%), respectively, suggesting that cell depolarization leads to a significant and specific change in the gene binding profiles of α1CCT, α1ACT, and α1HCT (Figure 4B).

**Figure 4.**
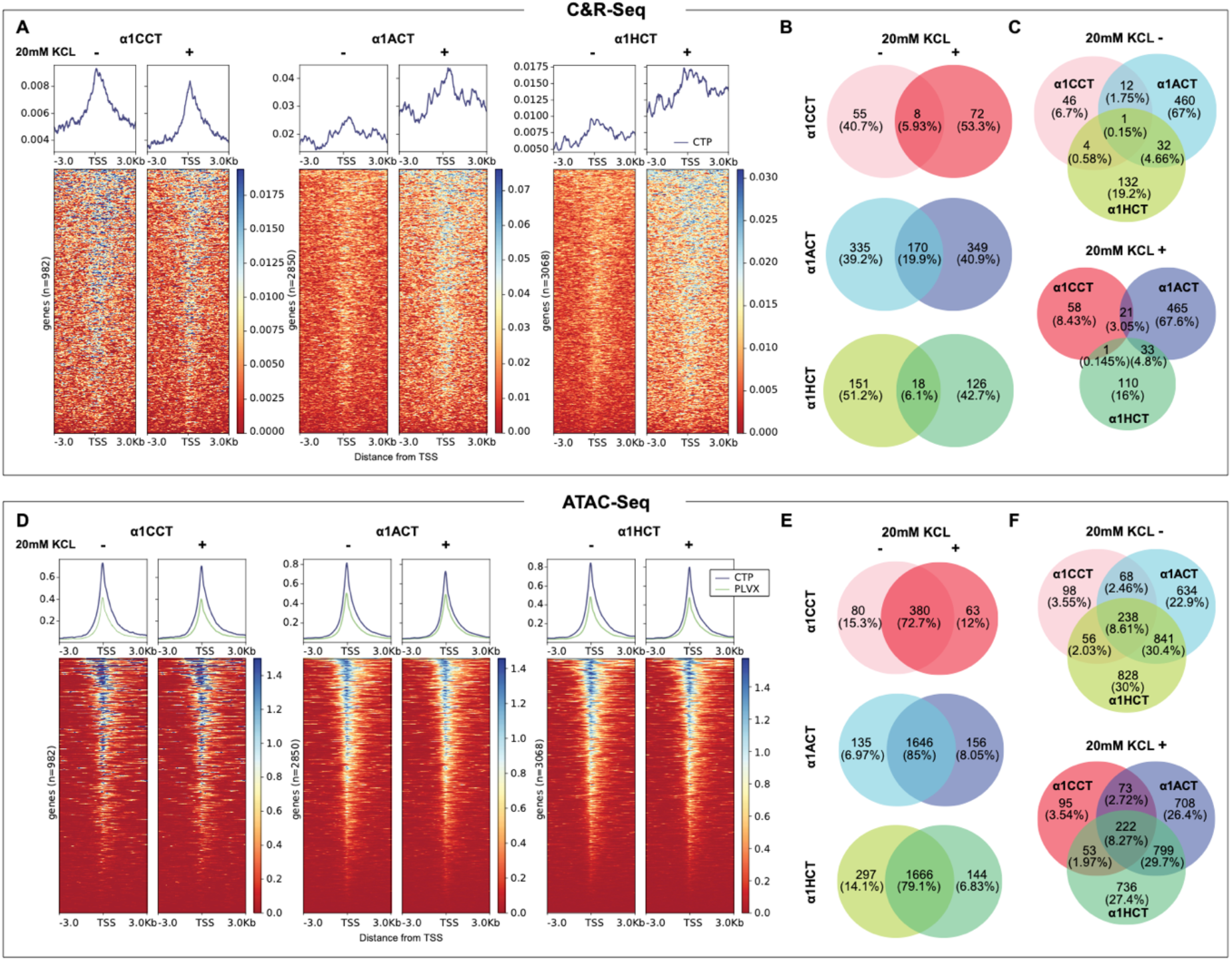
*α*1CCT, *α*1ACT, and *α*1HCT bind to genomic DNA in a Ca*²*⁺-dependent manner and alter chromatin accessibility independently of Ca*²*⁺ influx. See also Figure S4. (A) α1CCT, α1ACT, and α1HCT putative DNA binding profile within **±**3000bp relative to the transcription start site (TSS) of DEGs in both resting and depolarized cells. Heatmaps display the CUT&RUN signal intensity across target sites, with corresponding aggregate signal profiles (above). The intensity score was derived from the sequencing read coverage. (B) Venn diagrams indicate the limited overlaps in DEGs-associated genomic binding sites in resting and depolarized cells for α1CCT, α1ACT, and α1HCT. (C) Venn diagrams indicate overlaps in DEGs-associated genomic binding sites for α1CCT, α1ACT, and α1HCT in resting and depolarized cells, highlighting the unique binding patterns among the three CTPs. (D) α1CCT, α1ACT, and α1HCT affects chromatin accessibility distribution **±**3000bp TSS in resting and depolarized cells. Heatmaps show chromatin accessibility as measured by ATAC-seq, with aggregate signal profiles, pLVX is control, shown in green (above). (E) Venn diagrams indicate the prominent overlaps in chromatin accessibility of DEG-regulatory regions between resting and depolarized cells induced by α1CCT, α1ACT, and α1HCT. (F) Venn diagrams illustrate the overlap in chromatin accessibility of chromatin-accessible DEG- regulatory regions among α1CCT, α1ACT, and α1HCT in resting and depolarized cells. Shared and unique regions emphasize their differential impacts on chromatin structure.

Collectively, these observations indicate that α1CCT, α1ACT, and α1HCT exhibit physiological DNA binding properties near TSS regions and have distinct binding profiles. Moreover, these profiles are significantly altered under cell-depolarizing conditions. Our findings strongly support the notion that all three CTPs function as classical TFs and are involved in the Ca^2+^-dependent transcriptional regulation of specific target genes in hNPCs.

### VGCC CTPs increase chromatin accessibility in hNPCs

To assess potential global transcriptional alterations mediated by α1CCT, α1ACT, and α1HCT, we probed the chromatin accessibility changes, using ATAC-seq in the stable hNPC lines expressing either α1CCT, α1ACT, α1HCT, or empty pLXV vector as control. The specific chromatin binding sites associated with α1CCT, α1ACT, and α1HCT were identified by comparing the patterns of accessibility seen in the presence and absence of CTPs. Given that nuclear translocation of α1CCT and α1ACT, and the gene binding patterns of all three CTPs, are affected by intracellular Ca^2+^ levels, we performed ATAC-seq on the hNPC lines in the presence and absence of 20mM KCl. All three CTPs modify chromatin accessibility within ±3000 bp of annotated TSS (Figure 4D). Surprisingly, cell depolarization had little effect on global chromatin alterations associated with these three CTPs, with 72.7%, 85%, 79.1 % and of ATAC-seq peaks overlapping in resting versus depolarized cells, for α1CCT, α1ACT, and α1HCT, respectively (Figure 4E). Furthermore, approximately 238 of ATAC-seq peaks overlap among these three CTPs in resting cells, and 222 in the presence of 20mM KCl (Figure 4F), suggesting that the CTPs have overlapping effects on global transcriptional alterations, despite their distinct gene binding profiles induced by cell depolarization.

### *α*1CCT, *α*1ACT, and *α*1HCT mediate a neurodevelopmental program in a calcium-dependent manner in hNPCs

To explore the functional pathways regulated by α1CCT, α1ACT, or α1HCT through chromatin modulation or binding, we conducted RNA-seq analyses on stable hNPC cell lines expressing α1CCT, α1ACT, or α1HCT, at DIV5, and compared them with the empty pLVX vector in resting conditions only. Using RNA-seq, we identified 1007, 2949, and 3192 differentially expressed genes (DEGs) in α1CCT-, α1ACT-, or α1HCT- overexpressing hNPCs, respectively (Figure 5A). Integration of the CUT&RUN-seq and RNA-seq analyses revealed that 13.4%, 29%, or 9.24% of RNA-seq DEGs were correlated with direct DNA binding for α1CCT, α1ACT, or α1HCT (C&R-DEGs), respectively (Figure S5A). Therefore, α1CCT, α1ACT, or α1HCT, as independent DNA binding proteins, target specific genes and directly regulate their expression.

**Figure 5.**
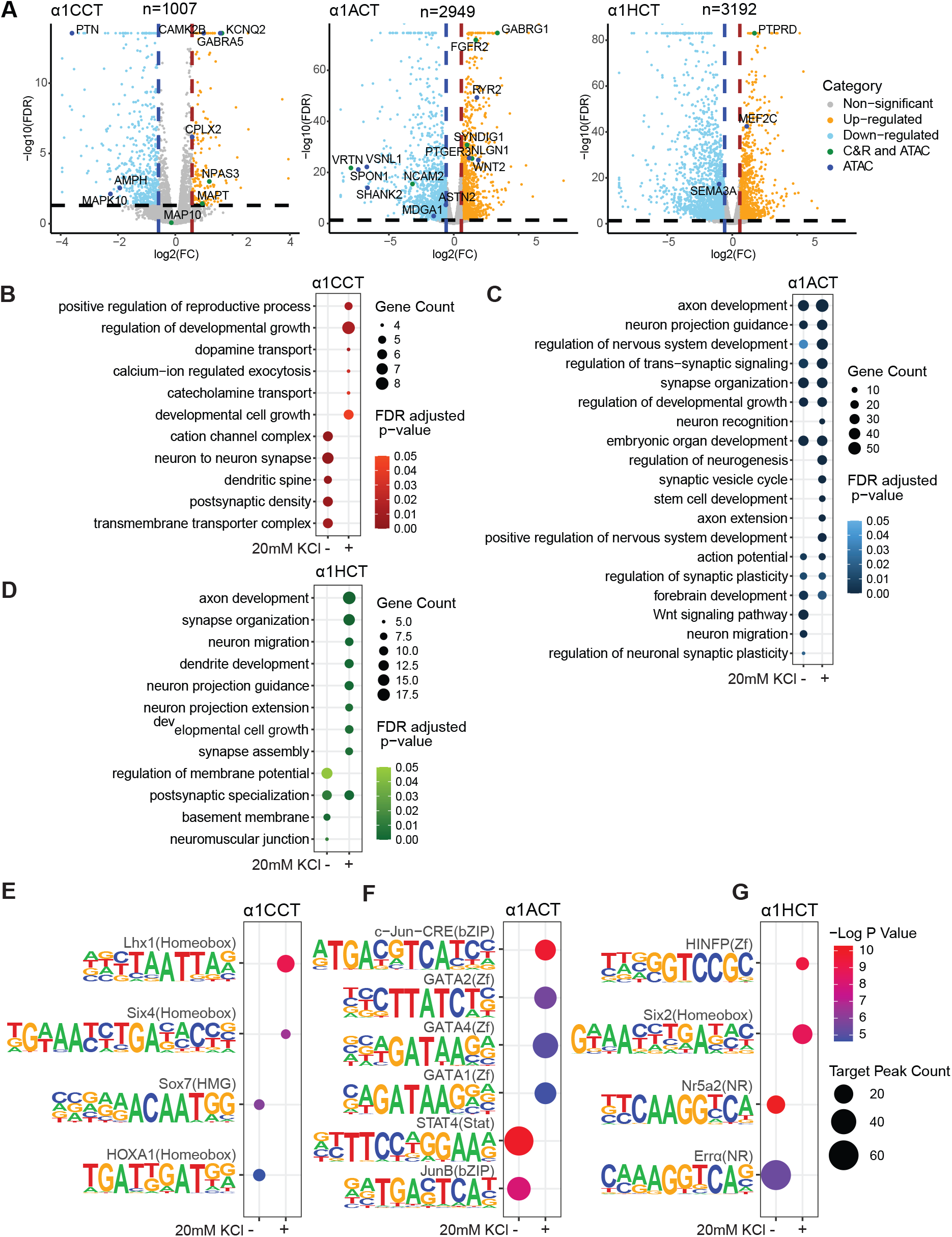
*α*1CCT, *α*1ACT, and *α*1HCT promote a genetic neural differentiation program in **hNPCs by binding to DNA in a Ca^2+^ dependent manner. See also Figure S5.** (A) Volcano plots of RNA-seq DEGs, the qRT-PCR-verified DEGs are highlighted as two categories, either with regulatory binding sites from both CUT&RUN-seq and ATAC-seq, or from ATAC-seq only. (B) Distinct enriched GO terms for RNA-seq DEGs directly regulated by α1CCT, inferred by CUT&RUN-seq, in hNPCs stably expressing α1CCT, with or without 20 mM KCl treatment. (C) Top enriched GO terms for RNA-seq DEGs directly regulated by α1ACT, inferred by CUT&RUN-seq, in hNPCs stably expressing α1ACT, with or without 20 mM KCl treatment. (D) Distinct enriched GO terms for RNA-seq DEGs directly regulated by α1HCT, inferred by CUT&RUN-seq, in hNPCs stably expressing α1HCT, with or without 20 mM KCl treatment. (E, F, and G) Distinct enriched DNA motifs from CUT&RUN-seq binding sites with an RNA- seq DEG annotation, for α1CCT (E), α1ACT (F), and α1HCT (G) with and without 20mM KCl treatment.

To correlate these results with those assessing chromatin accessibility, we integrated the ATAC-seq and RNA-seq analyses, which showed that 51.9%, 65.7%, or 66% of RNA-seq DEGs overlapped with ATAC-seq peaks (ATAC-DEGs) for α1CCT, α1ACT, or α1HCT, respectively (Figure S5B). These findings demonstrate that the three VGCC CTPs alter the transcriptional profile in hNPCs by modulating chromatin accessibility.

We performed Gene Ontology (GO) term analysis on RNA-seq integrated genes that were also identified by ATAC-seq or CUT&RUN-seq. For α1CCT, ATAC-DEGs showed enrichment in pathways related to synaptic transmission, axonogenesis, and transmembrane transport under both conditions (Figure S5C). However, α1CCT-associated C&R-DEGs were specifically linked to ion transmembrane transport and developmental growth, with these associations observed only under cell depolarizing conditions (Figure 5B). For α1ACT, ATAC- DEGs and C&R-DEGs were associated with neurogenesis, embryonic organ development, and synapse formation, in both high and low KCl conditions (Figure S5D). However, in the cell- depolarized condition, more neurogenesis-related genes and pathways were annotated among the DEG-C&R genes (Figure 5C). The results indicate that α1ACT regulates a set of neurogenesis genes basally but activates a larger ensemble for neurogenesis during neuronal activity. The top-ranked α1HCT ATAC-DEGs were associated with axon development, neuronal projection, development and guidance, and cell migration in both conditions (Figure S5E). For the α1HCT- associated C&R-DEGs, neuron-related pathways, including axon development, neuron migration, and synapse organization, were identified exclusively under cell-depolarized conditions (Figure 5D).

These results revealed that, among the DNA binding-regulated DEGs for α1CCT, α1ACT, and α1HCT, GO terms related to neuronal growth/neurogenesis, synapse formation, and ion channels are enriched, and overlapped. More interestingly, for α1CCT, α1ACT and α1HCT, neuronal growth-related pathways were predominantly enriched in C&R-DEGs in depolarized cells, a pattern not observed in ATAC-DEGs. These findings indicate that the transcriptional regulation of neuronal growth pathways associated with the three VGCC CTPs is enhanced by VGCC-induced Ca²⁺ signaling.

### *α*1CCT, *α*1ACT, and *α*1HCT have distinct calcium-responsive regulatory profiles in hNPCs

To further dissect the regulatory mechanisms underlying these transcriptional changes, we performed TF motif enrichment analysis using C&R-DEGs, which uncovered unique TF binding profiles for α1CCT, α1ACT, and α1HCT in hNPCs under basal and depolarized conditions. α1CCT exhibited a binding profile enriched in motifs for homeobox (HOXA1) and HMG (Sox7) TFs in basal conditions, while in depolarized states, it was associated with motifs for homeobox factors (Six4, Lhx1) (Figure 5E). α1ACT bound to motifs for bZIP (JunB), and STAT (STAT4) families in basal conditions, whereas in depolarized conditions, it recognized motif for bZIP(c-Jun-CRE) and ATAA/TATT (GATA 1.2.3.4) factors (Figure 5F). α1HCT had a binding profile enriched for nuclear receptor (NR) motifs (ERRα, Nr5a2) in basal conditions, while in depolarized states, it recognized Homeobox (Six2) and Zf (HINFP) motifs (Figure 5G). These motif binding profiles suggest that α1CCT, α1ACT, and α1HCT engage in different neuronal activity-related regulatory programs, underscoring their potential roles in neural differentiation and development. While their motif preferences are distinct, their biological functions appear to overlap.

Our findings indicate that the distinct transcriptional regulation mediated by α1CCT, α1ACT, and α1HCT is driven by their ability to interact with specific TF motifs, dynamically responding to KCl-induced depolarization. This coordination links their chromatin-modifying functions to transcription factor-driven gene expression changes, highlighting their role as Ca^2+^- dependent regulators of transcription in hNPCs.

### Verified DEGs for *α*1CCT, *α*1ACT, and *α*1HCT in neuronal expression and synaptic regulation

Following the identification of DEGs for α1CCT, α1ACT, and α1HCT through next-generation sequencing, we validated key targets using TaqMan qRT-PCR assays, focusing on genes that overlapped with CUT&RUN (C&R) and ATAC-seq data. The validated α1CCT target genes, including *KCNQ2*, *MAPT*, and *NPAS3*, are strongly associated with neuronal excitability and ion translation. Additionally, we verified ATAC-seq confirmed genes such as *PTN*, *MAPK10*, *GABRA5*, *CAMK2B*, and *CPLX2*, reinforcing the involvement of α1CCT in synaptic plasticity, neurotransmission, and Ca^2+^-dependent signaling (Figure S5F). α1ACT directly regulates genes such as *FGFR2*, *SYNDIG1*, *NCAM2*, *GABRG1*, and *NLGN*, indicating roles in neurogenesis, synapse formation, and synaptic adhesion (Figure S5G). Verification of ATAC-DEGs—*ASTN2*, *MDGA1*, *VSNL1*, *SPON1*, *WNT2*, *RYR2*, *SHANK2*, and *PTGER3*— suggests that α1ACT has additional functions in neuronal migration, Wnt signaling, and excitatory synapse maturation (Figure S5G). For α1HCT, validated genes, such as *PTPRD*, suggest roles in neuronal connectivity and synapse organization, while the ATAC-seq-confirmed targets *SEMA3A* and *MEF2C* highlight its function in axon guidance, neuronal migration, and synaptic remodeling (Figure S5H). This suggests that α1HCT preferentially regulates axon development and neural circuit formation. These validated genes highlight that each VGCC CTP regulates distinct, yet overlapping, transcriptional programs, contributing to neuronal differentiation, synaptic organization, and neurodevelopmental signaling pathways.

### *α*1CCT, *α*1ACT, and *α*1HCT induce a neuronal phenotype in hNPCs

The transcriptional changes induced by α1CCT, α1ACT, and α1HCT suggest a broader impact on neuronal differentiation and morphology. Consistent with these findings, we observed that many of the validated DEGs are linked to neurite outgrowth, synaptic development, and cytoskeletal dynamics, raising the possibility that these VGCC CTPs may promote structural changes in hNPCs. While conducting genomic and transcriptomic studies on hNPCs stably expressing CTPs, we observed notable changes in cellular phenotype, with increases in neurite morphologies. We found that hNPCs expressing any one of the three CTPs grew significantly more neurites than the empty vector control stable line at DIV5 (α1CCT: 2.90±0.22 neurites; α1ACT: 2.75±0.23 neurites; α1HCT: 3.63±0.25 neurites vs control: 1.53±0.16 neurites, p < 0.002) (Figure 6A-B). Additionally, hNPCs expressing α1CCT, α1ACT and α1HCT significantly increased the “longest neurite length” compared to the control stable line (α1CCT: 90.75±10.60 μm; α1ACT: 66.55±7.68 μm; α1HCT: 74.62±9.16 μm, p < 0.001vs control: 32.44±5.46 μm) and the average neurite length (Figure 6C and D).

**Figure 6.**
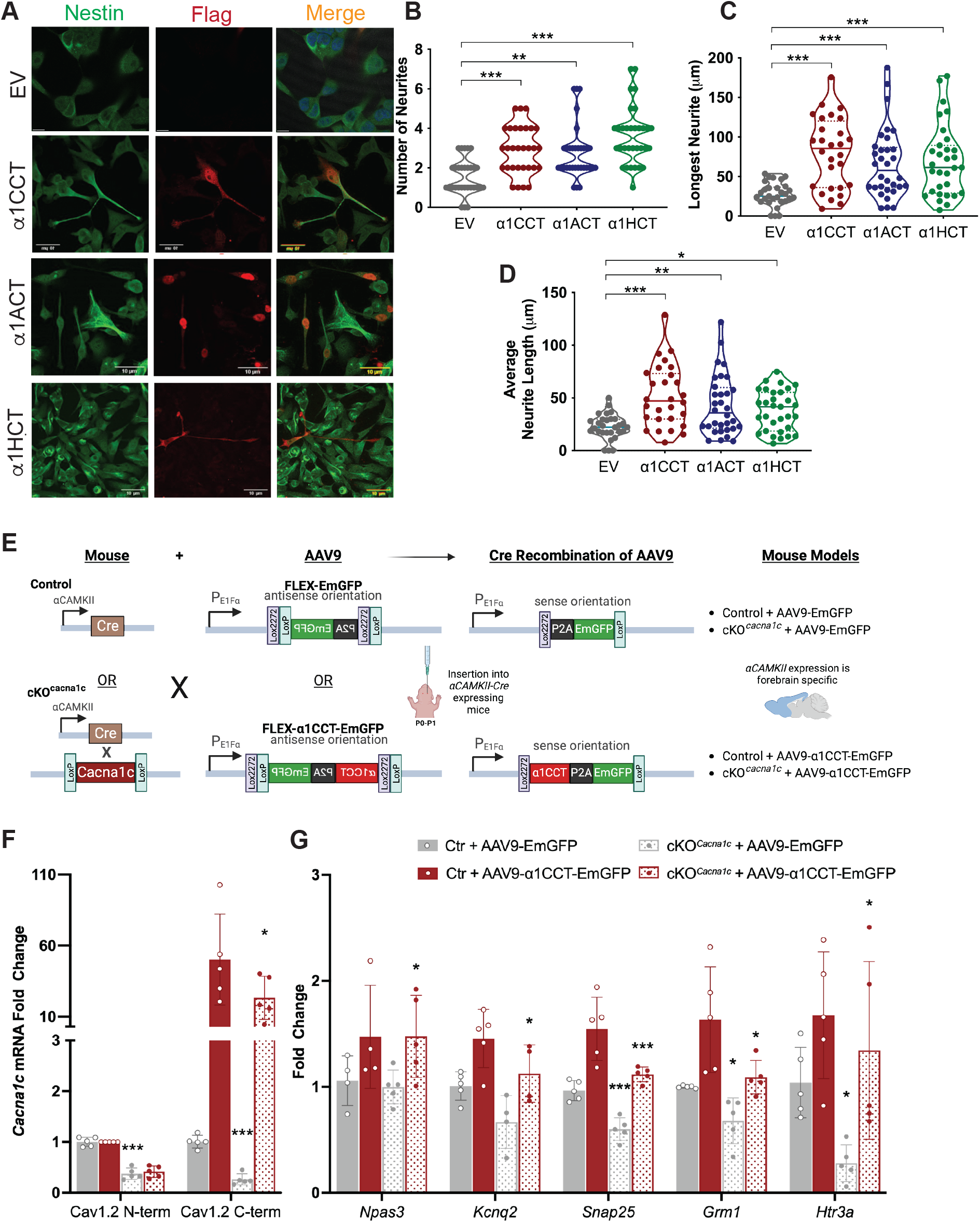
*α*1CCT, *α*1ACT, and *α*1HCT promote neurite outgrowth in hNPCs, and *α*1CCT restores expression of critical neuronal genes in a conditional forebrain *Cacna1c* knockout mouse model. (A) Representative images of hNPCs stably expressing C-terminal 3XFLAG tag-labeled α1CCT, α1ACT, or α1HCT co-stained with antibodies directed against the hNPC marker Nestin (green) or 3XFLAG (red). (B, C, and D) Quantification of neurites per cell, longest neurite per cell, and average neurite length (n > 30 cells, *p<0.05, **p<0.01, ***p<0.001). (E) Schematic diagram showing the AAV-FLEX-EmGFP or AAV-FLEX-α1CCT-EmGFP viral vectors used for *in vivo* injections and the derived mouse models. (F) Validation qRT-PCR analysis of *Cacna1c* knockout and α1CCT re-expression using primers directed at the 5’ or 3’ ends of the gene, respectively. (G) qRT-PCR analysis of selected upregulated and downregulated DEGs identified through RNA-seq in microdissected mouse forebrain tissue N = 4-5 mice per condition, *p<0.05, **p<0.01, ***p<0.001

Taken together, these findings indicate that the novel VGCC CTPs, α1CCT, α1ACT, and α1HCT mediate widespread changes in gene expression, promoting the development or maturation of a neuronal phenotype.

### *α*1CCT restores disrupted gene expression in a forebrain-specific *CACNA1C* knockout mouse

We previously demonstrated that transgenic expression of α1ACT in *CACNA1A*-deficient mice improves survival and ataxic behavior, and restores expression of α1ACT-regulated genes (Du et al., 2019). L-type VGCCs, and most notably Ca_v_1.2 (encoded by *Cacna1c*), play a critical role in both cardiac and neuronal function, as well as activity Ca^2+^-dependent transcriptional regulation (Dolmetsch et al., 2001; Kandel, 2001; Li et al., 2016a; Raymond, 2007; Wheeler et al., 2008; Wheeler et al., 2012; Wu et al., 2001). As a test case to relate CTP-regulated gene expression to the parent VGCC gene *in vivo*, we used AAV9-mediated delivery of α1CCT in a conditional forebrain knockout mouse model (Forebrain*-Cacna1c* cKO), as complete *Cacna1c* knockouts are non-viable. In this mouse model, a floxed *Cacna1c* allele is deleted selectively in forebrain neurons expressing Cre under a CaMKII transgene promoter (Figure 6E and S6A) (Lee et al., 2012).

To selectively express α1CCT in Cre-expressing, *Cacna1c*-null neurons, we employed AAV9 vectors that use a Double Flox Inverted Open Read Frame (DIO) system, enabling conditional expression of α1CCT protein in forebrain neurons. Forebrain-*Cacna1c* cKO mice (P0-P1) were injected via the lateral ventricles with AAV virus expressing either control EmGFP (AAV9-FLEX-EmGFP) or α1CCT (AAV9-FLEX- α1CCT:P2A:EmGFP) (Figure 6E and G). At P30-35, forebrain tissue was sectioned for laser capture microdissection (LCM) to isolate EmGFP-labeled cortical neurons for TaqMan qRT-PCR analysis (Figure 6F and G).

Expression analysis revealed significant downregulation of α1CCT-regulated DEGs, such as *Npas3* (-11.54±7.7%)*, Kcnq2* (-24.79±12.0%)*, Snap25* (-36.41±8.8%)*, Grm1*(-32.03±10.0%), and *Htr3a* (-76.1±17.1%) in the Forebrain-*Cacna1c* cKO mice receiving AAV-EmGFP virus, compared to wild-type mice receiving AAV-EmGFP virus (Figure 6G). More importantly, the expression of these genes was rescued to normal or significantly higher levels in cells expressing α1CCT (*Npas3:* 47.64±12.0%; *Snap25:* 51.7±6.7%; *Grm1*(41.13±13.6%)*, Htr3a:* 106.43±30.7%) (Figure 6G).

These results show that lack of *Cacna1c* impacts the gene expression profile of α1CCT in cortical neurons, but *in vivo* re-expression of α1CCT alone rescues the expression of both α1CCT-regulated DEGs and chromatin accessibility DEGs. These rescued genes are involved in synaptic transmission (*Snap25*), ion transport (*Kcnq2*), neurogenesis (*Npa3*), and neurotransmitter receptors (*Grm1*and *Htr3a*). Taken together, these findings indicate that α1CCT, as an independent protein, plays an important role in promoting maturation of a neuronal phenotype.

## Discussion

This study provides evidence for the presence of a conserved genetic strategy across the entire VGCC gene family that enables direct signaling to the nucleus by a set of C-terminal transcription factors coupled to, and co-expressed with, the VGCC channel proteins. The co-expressed C-terminal proteins are encoded by overlapping cistrons in the same mRNA. Translation of the 70-75kD CTPs, α1CCT, α1ACT, and α1HCT, occurs by means of IRES-like structures within the α1C, α1A, and α1H mRNAs. In addition to the *genetic* coupling of the channel protein to the co-expressed transcription factor, we have identified *physiological* coupling of ion channel activity to the behavior of the same transcription factor. We find that the nuclear translocation of α1CCT and α1ACT is influenced by cellular Ca^2+^ currents in neurons in real-time. Moreover, cell-depolarizing conditions dramatically alter the gene binding profiles of α1CCT, α1ACT and α1HCT, indicating a *direct link* between Ca²⁺-dependent VGCC activity and transcriptional regulation. These Ca^2+^ currents facilitate the nuclear-cytoplasmic translocation of VGCC CTPs, which in turn directly activates ensembles of DEGs that are crucial for the development of neuronal phenotypes, particularly neurogenesis, synaptic formation, and ion channel transport, in concert with the emergence of neuronal activity.

Ultimately, the transcriptional functions of α1CCT, α1ACT and α1HCT are highly integrated with Ca²⁺-induced nuclear signaling, shaping neuronal connectivity and synaptic function.

Co-expression of membrane receptors and nuclear signaling molecules has been typically associated with a protease cleavage event that liberates the signaling molecule, as in the cases of amyloid precursor protein, Notch receptor, and ERBB2/Her2, although the capacity for IRES-based expression has also been reported in some isoforms of Notch receptor and ERBB2/Her2 (Godfrey et al., 2022; Karginov et al., 2017; Lauring and Overbaugh, 2000). While our findings do not exclude the existence of other cellular pathways for generating Ca^2+^ channel CTPs, such as protease cleavage and cryptic promoters (Gomez-Ospina et al., 2013; Hulme et al., 2006), this study demonstrates the pervasive expression of CTPs broadly within the VGCC gene family by an internal translation process. Of these, we formally confirmed that α1CCT, α1ACT, and α1HCT, representing the three subfamilies of VGCC genes, are translated by a cap-independent translation mechanism. Moreover, this expression strategy of CTPs appears to be accompanied by nuclear signaling, as we identified CTPs expressed by all ten VGCC genes translocating to the nucleus.

Key domains within the Ca^2+^ channel C-termini have been identified as mediators of Ca^2+^ channel function, specifically those that competitively modulate Ca^2+^ channel gating processes by inhibiting Ca^2+^-dependent inactivation (CDI) (Bock et al., 2011; Castiglioni et al., 2006; Liu et al., 2017; Singh et al., 2006; Wahl-Schott et al., 2006), or that bind synaptic vesicles to initiate their delivery to the active zone (Gardezi et al., 2016; Wong et al., 2014). In some of these studies, the capacity for modulation of full-length Ca^2+^ channel function via the C-terminus was noted even when C-terminal fragments were co-expressed with full-length channels. Thus, we cannot exclude an additional role for cytoplasmic CTPs in potentially modulating Ca^2+^ currents to influence their nuclear translocation. As shown here, all ten VGCC CTPs distribute between the cytoplasm and nucleus, albeit at different rates, and translocation of α1CCT and α1ACT was driven by Ca^2+^ influx through specific channels. Interestingly, none of the brief stimuli used in this study had a significant effect on the subcellular localization of α1HCT, even though there was a trend towards increasing nuclear Ca^2+^ fluorescence in live primary rat cortical neurons. It is possible that the localization and translocation of the *CACNA1H*-derived CTP may be independent of Ca^2+^signaling over the short time course used here, or through other processes.

Ca^2+^ is a ubiquitous cellular second messenger that plays a vital role in almost every facet of neuronal development, maturation, and maintenance, by signaling to TFs, such as NFAT and CREB (Hardingham et al., 2001; Li et al., 2016b; Wheeler et al., 2012). VGCC channels link Ca^2+^ entry through electrical activity to nuclear gene expression. VGCCs serve as key Ca^2+^ signal transducers, and Ca^2+^ influx through them mediates a wide variety of downstream physiological events (Berridge, 2014). Previously, Gomez-Ospina et al. showed that the CTP of Ca_V_1.2 (α1CCT) acts as an independent TF that regulates a group of neuronal genes, and that its translocation is influenced by Ca^2+^, although they did not elucidate the relationship between Ca^2+^ signaling and gene regulation. Yang et al. reported the role of cytosolic and nuclear DCT of Ca_V_1.3 (α1DCT) in regulating neurogenesis and channel activity. Our group previously demonstrated that α1ACT is a critial TF in cerebellar development (Du et al., 2013; Du et al., 2019; Gomez-Ospina et al., 2006; Yang et al., 2022). This study extends the findings of previous Ca^2+^ channel studies and paints a global picture of Ca^2+^ channel direct signaling to the nucleus, by establishing an additional connection between Ca^2+^ and the secondary CTP gene products of the VGCC genes.

In this study, we utilized the CUT & RUN-seq technique, coupled with ATAC-seq and RNA-seq, to analyze the genomic and transcriptomic profiles of α1CCT, α1ACT, and α1HCT expressed in hNPCs. Our findings reveal that, for all three CTPs, Ca^2+^ influx plays a critical role in mediating their biological function. On the one hand, 222 (resting)/238 (depolarized) genes that show altered chromatin accessibility have been identified in common among three CTP- overexpressing hNPCs. Interestingly, the majority of these gene appear to function in promoting neuronal growth pathways and are not influenced by Ca^2+^ influx. On the other hand, each CTP is associated with distinct TF motifs, without overlap among all three, and this could contribute to the regulation of specific genes for neurodevelopment, synaptic formation, and ion membrane transport in response to Ca^2+^ influx. For instance, an AT-rich motif was previously identified for α1ACT (Du et al., 2013). Meanwhile, there was a 20 - 30% overlap between DEGs in hNPCs and those associated with neurite outgrowth, with 10 of these genes verified in α1ACT- expressing transgenic mice and rescued by α1ACT (Du et al., 2019). These findings reinforce the established role of Ca^2+^ as a second messenger, by demonstrating that the three co-expressed VGCC proteins, α1CCT, α1ACT and α1HCT, drive expression of neuronal genes. These CTPs act in concert with the emergence of neuronal activity, enhancing Ca^2+^ efficiency as a mediator of nuclear-cytoplasmic translocation and gene regulation.

While the genetic organization observed for the Ca^2+^ channel and its CTPs appears to facilitate efficient coordination of gene activation, it also increases the likelihood that genetic disruptions will have intricate downstream consequences and may help explain some of the complex clinical phenotypes seen with VGCC gene mutations. Loss- or gain-of-function mutations in the *CACNA1C* gene, encoding the α1C (Ca_v_1.2) L-type Ca^2+^ channel subunit, are associated with a pleiotropic set of disorders including the autism spectrum disorder (ASD) Timothy Syndrome, bipolar disorder, schizophrenia, and major depressive disorder (MDD), as well as severe cardiac defects (Boczek et al., 2013; Boczek et al., 2015; Cross-Disorder Group of the Psychiatric Genomics, 2013; Dedic et al., 2018; Green et al., 2010; Splawski et al., 2004).

Loss- or gain-of-function mutations in the *CACNA1A* gene, encoding the α1A (Ca_v_2.1) P/Q-type Ca^2+^ channel subunit, may lead to ataxia, hemiplegic migraine, epilepsy, or cognitive disability (Indelicato et al., 2018). Mutations in the α1H (Ca_v_3.2) subunit, part of the T-type VGCC, are associated with chronic neuropathic pain disorders, ASD, and several types of mental illness (Becker et al., 2017; Carter et al., 2019; Heron et al., 2007; Lee et al., 2018; Rzhepetskyy et al., 2016; Souza et al., 2016; Splawski et al., 2006; Stringer et al., 2020). Given that such mutations across the coding regions of these genes may have a positive or negative effect on the function of a channel, the co-expressed transcription factor, or the regulation or translocation of that transcription factor, this diversity of phenotypes may arise from a disruption of the complex dynamics of these two-protein/one-gene systems. Unsurprisingly, attempts to recapitulate such phenotypically complex disorders in animal models have often fallen short of providing an understanding of pathogenesis.

In conclusion, this study identifies a pervasive dual protein expression system within the VGCC gene family and sheds new light on the intricate genetic and physiological coordination inherent in members of the VGCC gene family. Our findings establish a direct link between Ca^2+^ channel activity and nuclear gene expression through the co-expression and translocation of CTPs. This dual-protein mechanism underscores the importance of Ca^2+^ as a critical second messenger that extends its influence from the cellular membrane, where Ca^2+^ channels reside, to the nuclear CTPs, thereby orchestrating complex gene regulatory networks essential for neurogenesis, synaptic formation, and ion channel transport. This coordinated process adds to the previously identified TFs a new set of gene regulatory proteins, the *calcium channel-coupled transcription factors*, as CTPs co-expressed with the VGCC channel proteins. The operation of this dual-protein system as a self-regulatory mechanism under physiological conditions will require further study. Additionally, since mutations in these genes result in multifaceted clinical phenotypes, this study provides valuable insights into the mechanisms that underlie neuronal phenotypes and their potential perturbations in disease states, paving the way for novel therapeutic strategies to target the VGCC gene family, such as modulating channel function, RNA silencing, or viral delivery of CTP transcription factors.

## Statistical analysis

Statistical analyses were conducted using GraphPad Prism 7 software (GraphPad Software Inc., La Jolla, CA), RStudio (Boston, MA), and SPSS (IBM, Armonk, NY). For single comparisons, Student’s t test was used to establish statistical significance. For multiple sample comparisons, One-Way ANOVA was performed, followed by Student-Newman-Keuls Method post hoc tests. Data were expressed as the mean ± SEM and written with the identification of n under appropriate figure legends, with *p < 0.05, **p < 0.01; ***p < 0.001.

## Data and Code availability

All data reported in this paper will be publicly available upon manuscript publication in an online database or via request to the corresponding authors. All source code will be deposited at Github. Further additional information and requests should be addressed to lead contact, Christopher M. Gomez

## Acknowledgments

This work was supported by NIH grants: R01NS082788, R01NS094665, R35NS116868 (C.M.G.), a Floyd family donation, and Margaret Hackett Family Program Translational Research Award. We thank Dr. Sally Rowland and Dr. Xiaoxi Zhuang for comments on the manuscript; Drs. Vytas Bindokas and Christine Labno at the University of Chicago Microscopy Core Facility for the imaging service.

## Author Contributions

E.R., X.D. and C.M.G. conceived the project. E.R., X.D. and C.M.G. designed the study and experiments. E.R., T.T., J.G., D.H., and E.G., performed the most experiments, Q.L. Y.L. and C.W. analyzed the ChIP and RNA-seq data. C.W. was in charge of mice breeding and phenotyping. E.R., T.T., X.D., and C.M.G. wrote the manuscript with input from all coauthors.

