## Supplemental Documents for "Calcium channel-coupled transcription factors facilitate direct nuclear signaling"

Inventory for Supplemental Information

A. **Supplemental Figure and Figure Legends 1 to 5**
B. **Supplemental Table 1 to 5**
C. **Supplemental Videos**

- **Video S1** Live cell imaging of EV-emGFP in cultured rat cortical neurons. Neurons were imaged for 10 minutes following glutamate uncaging.
- **Video S2** Live cell imaging of α1CCT in cultured rat cortical neurons. Neurons were imaged for 10 minutes following glutamate uncaging.
- **Video S3** Live cell imaging of α1ACT in cultured rat cortical neurons. Neurons were imaged for 10 minutes following glutamate uncaging.
- **Video S4** Live cell imaging of α1HCT in cultured rat cortical neurons. Neurons were imaged for 10 minutes following glutamate uncaging.

**D. Movie Legends**

Videos showing live cell imaging of cultured rat cortical neurons infected with AAV virus expressing either EV-emGFP (A), α1CCT-EmGFP (B), α1ACT-EmGFP (C), or α1HCT (D). Neurons were imaged for 10 minutes following glutamate uncaging. EV-emGFP showed no cyto-nuclear translocation, while α1CCT and α1ACT showing increased or decreased nuclear translocation following glutamate uncaging, respectively. Additionally, α1HCT showed a slight trend of increased nuclear translocation following uncaging. Scale bars represent 20 uM.

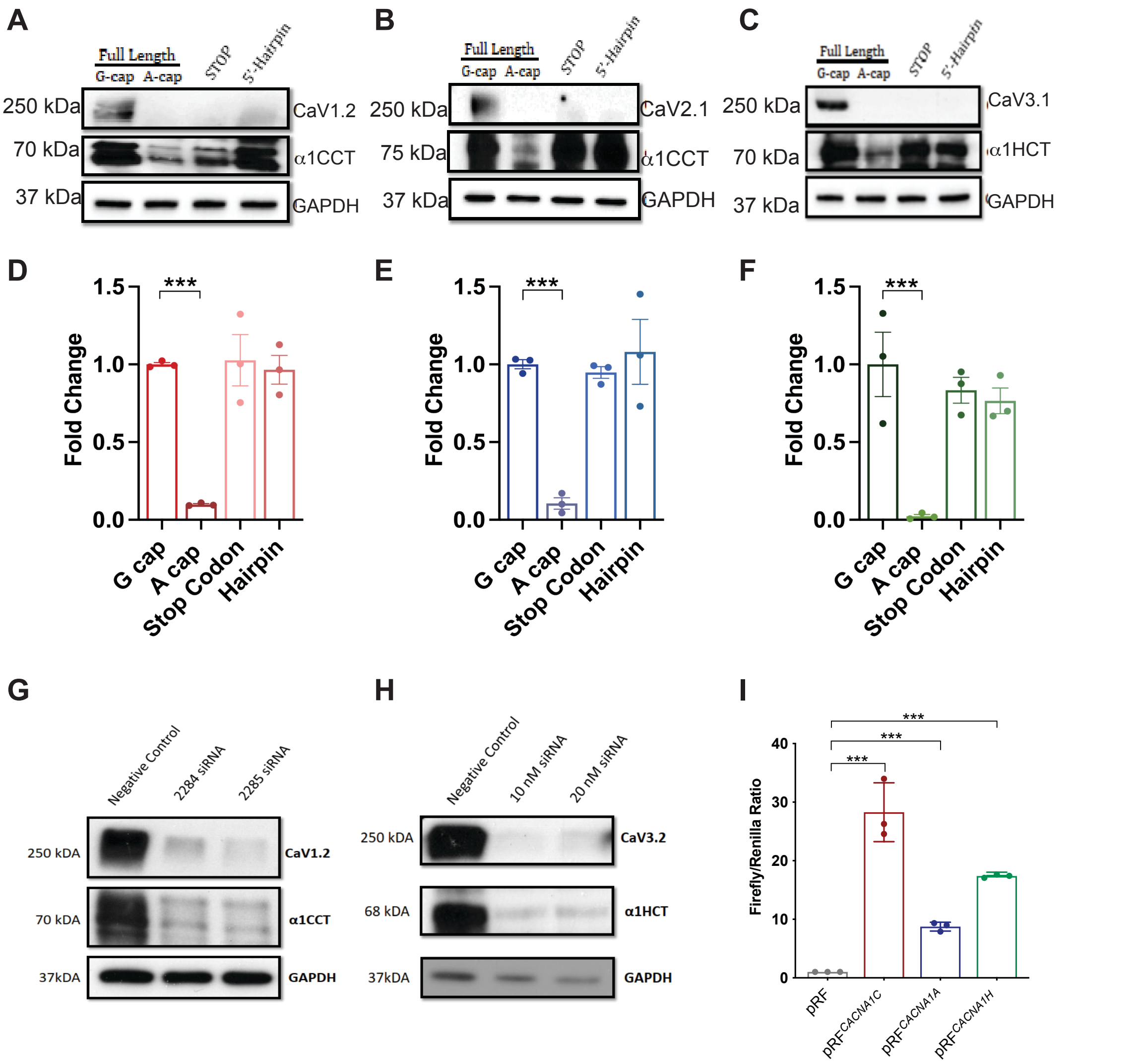


**Supplemental Figure 1. *CACNA1C, CACNA1A,* and *CACNA1H* mRNAs encode two distinct proteins from overlapping cistrons through a cap-independent mechanism.**

(A, B, and C) Western blot analysis of protein lysates from HEK293Tcells transiently transfected with *CACNA1C* (A), *CACNA1A* (B), or *CACNA1H* (C) *in vitro* transcribed mRNA. Full-length mRNAs were capped with either an m7G or an m7A cap. STOP constructs had one or two premature termination codons inserted upstream of C-terminal protein start sites. 5'-hairpin constructs had a large hairpin structure inserted directly downstream of the initiating methionine.

(D, E, and F) qPCR analysis of RNA collected from HEK293T cells transiently transfected with *CACNA1C* (D), *CACNA1A* (E), or *CACNA1H* (F) mRNA. (N = 6 for each condition).

(G, H) Western blot analysis of protein lysates from HEK293 of cell lines stably expressing either *CACNA1C* or *CACNA1H* and transfected with siRNAs directed towards the 5' ends of the *CACNA1C* or *CACNA1H* genes.

(I) Luciferase activity as measured by Firefly/Renilla ratio for the bicistronic vector pRF with 1000-bp insertions directly upstream from α1CCT, α1ACT, or α1HCT initiating methionines, compared to empty vector. (N = 3 for each condition, $p<0.001$).

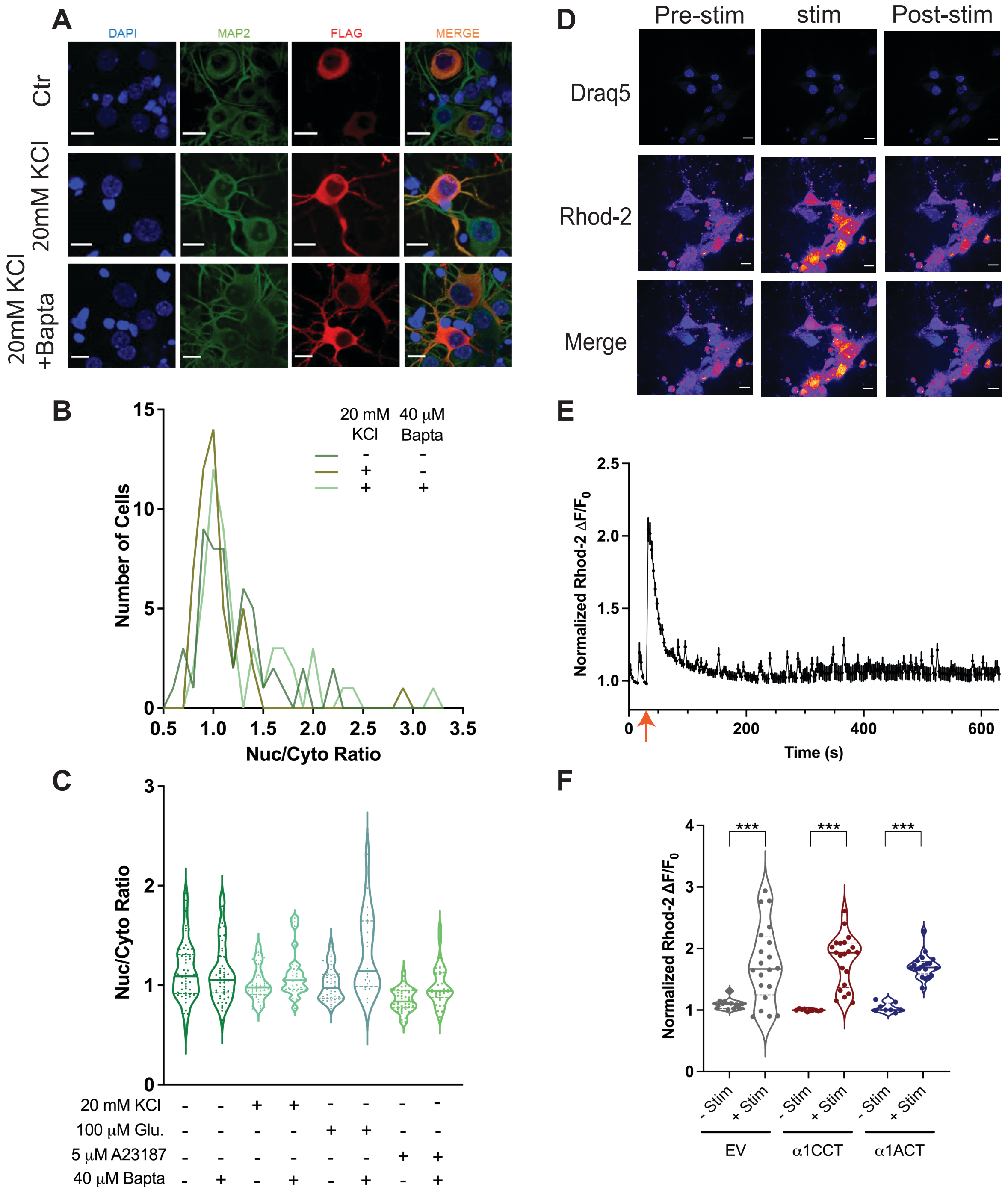


**Supplemental Figure 2. Neuronal depolarization does not shift α1HCT localization in fixed rat cortical neurons, and intracellular Ca2+ changes with uncaged glutamate stimulation of live rat cortical neurons expressing EmGFP, α1CCT, or α1ACT.**

(A) Representative images of fixed rat cortical neurons transfected with α1HCT mRNA and treated with 100 µM glutamate with or without BAPTA-AM. Fixed neurons were stained with DAPI (blue), MAP2 antibody (green), and 3XFLAG antibody (red). Scale bars = 10 microns.

(B) Quantification of nuclear/cytosolic fluorescence signal of rat cortical neurons transfected with α1HCT mRNA with 20mM KCl with or without BAPTA-AM.

(C) Quantification of nuclear/cytosolic fluorescence signal of rat cortical neurons transfected with α1HCT mRNA with different treatments.

Neurons were treated with either 20 mM $K^+$, 100 µM glutamate, or the calcium ionophore A23187 with or without a 5-minute pretreatment of BAPTA-AM.

N > 50 cells for each condition, *p<0.05, **p<0.01, ***p<0.001.

(D) Representative images of live rat cortical neurons loaded with both the live-cell nuclear stain Draq5 and the ratiometric calcium indicator Rhod-2, imaged pre-stimulation via uncaging glutamate (left panels), immediately after glutamate uncaging and consequent neuronal calcium stimulation (middle panels), and ten minutes post-stimulation (right panels). Scale bars = 10 microns.

(E) Representative imaging trace of Rhod-2 $\Delta F/F_0$ over a ten-minute imaging period in a live rat cortical neuron. Arrow indicates glutamate uncaging pulses.

(F) Quantification of Rhod-2 $\Delta F/F_0$ immediately pre- and post-stimulation in live rat cortical neurons.

N > 20 cells per condition for + Stim., N > 10 cells per condition for -Stim.

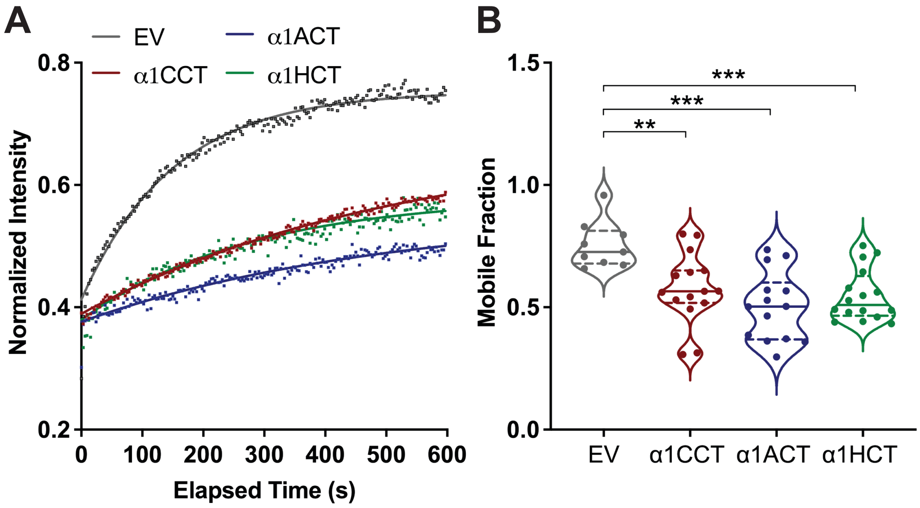


**Supplemental Figure 3. FRAP analysis of α1CCT, α1ACT, and α1HCT in cultured rat cortical neurons.**

(A) FRAP recovery curves showing the normalized intensity over time for cultured rat cortical neurons expressing EmGFP, α1CCT-EmGFP, α1ACT-EmGFP, or α1HCT-EmGFP. EV-expressing cells exhibit the highest fluorescence recovery, indicating greater mobility compared to α1CCT, α1ACT, and α1HCT. The recovery profiles represent a nonlinear fit to the average of individually photobleached cells imaged for 10 minutes post-bleach.

(B) Quantification of the mobile fraction from the FRAP analysis. Cells expressing α1CCT, α1ACT, and α1HCT all show significantly reduced mobile fractions compared to EV, indicating restricted mobility.

N > 10 cells per condition, *$p<0.05$, **$p<0.01$, ***$p<0.001$

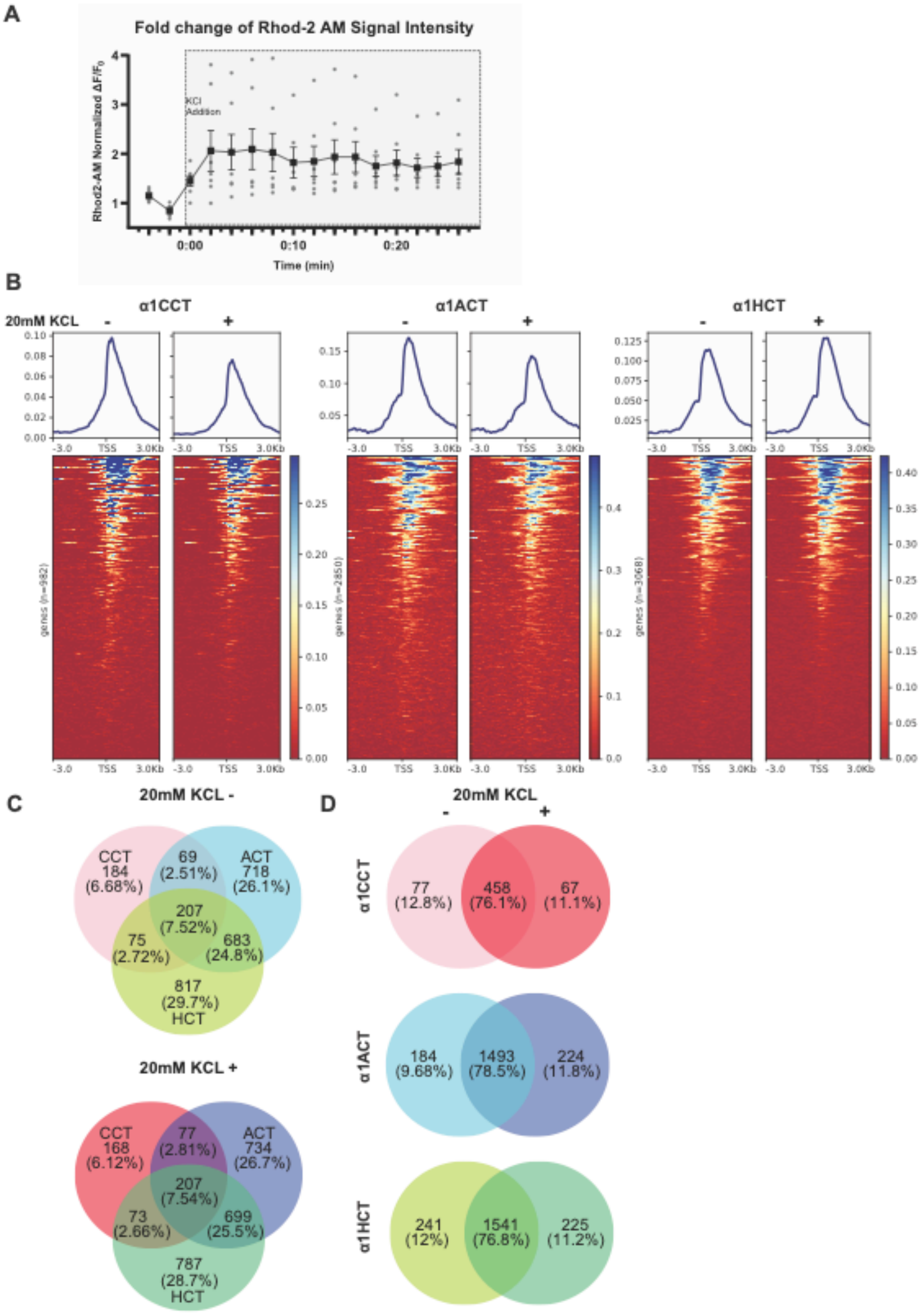


**Supplemental Figure 4. Intracellular Ca2+ changes with 20mM KCl, and CTPs' influence on the effects of H3K4me3-related genomic binding sites.**

(A) Imaging trace of Rhod-2 $\Delta F/F_0$ in human neural progenitor cells (n=7) over a 26-minute imaging period. 20mM KCl was added to media 10 seconds before 0:00 minutes after a baseline was established. KCl addition indicated by gray box.

(B) CUT&RUN-seq profiles for H3K4me3 enrichment distribution within ±3000bp TSS in hNPC stable cell lines expressing α1CCT, α1ACT, and α1HCT under resting and depolarized conditions (± 20 mM KCl).

(C) Venn diagrams depicting differentially enriched H3K4me3-associated DEGs in resting or depolarized conditions in hNPC stable cell linesexpressing α1CCT, α1ACT, and α1HCT. Numbers indicate the count of unique and shared H3K4me3-associated DEGs.

(D) Comparative Venn diagrams of H3K4me3-associated DEGs across hNPC stable cell linesexpressing α1CCT, α1ACT, and α1HCT in resting and depolarized conditions. The overlap between CTPs highlights distinct and shared regulatory elements modulated by H3K4me3.

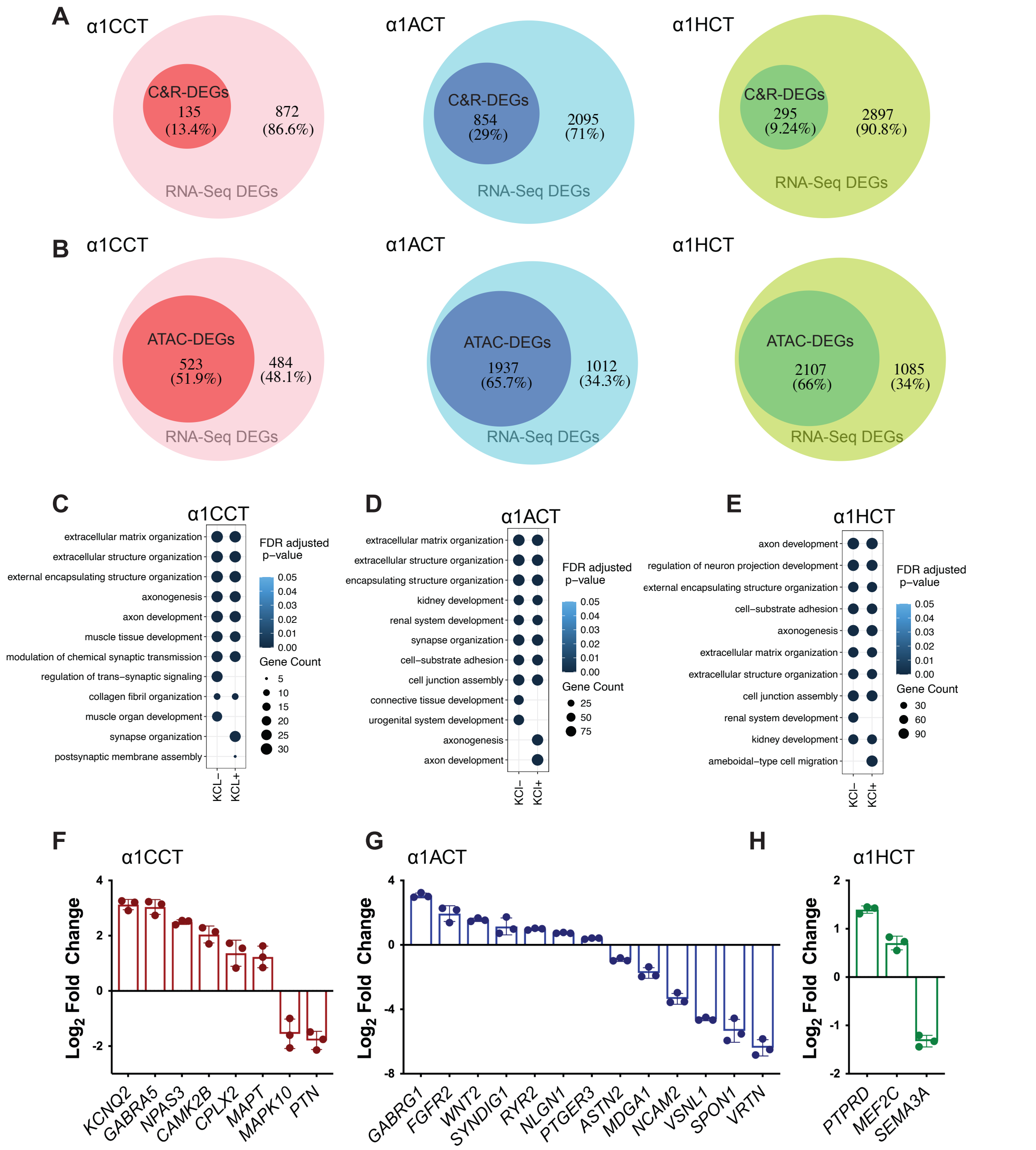


**Supplemental Figure 5. Integration of CUT&RUN-seq or ATAC-seq with RNA-seq in hNPCs stably expressing α1CCT, α1ACT, and α1HCT.**

(A) Venn diagrams showing the percentage of C&R -DEGs within RNA-seq-DEGs for α1CCT (left), α1ACT (middle), and α1HCT (right).

(B) Venn diagrams illustrating the percentage of ATAC-DEGs within RNA-seq-DEGs for α1CCT (left), α1ACT (middle), and α1HCT (right).

(C) Distinct enriched GO terms for RNA-seq DEGs directly regulated by α1CCT, inferred by ATAC-seq, in hNPCs stably expressing α1CCT, with or without 20 mM KCl treatment.

(D) Top enriched GO terms for RNA-seq DEGs directly regulated by α1ACT, inferred by ATAC-seq, in hNPCs stably expressing α1ACT, with or without 20 mM KCl treatment.

(E) Distinct enriched GO terms for RNA-seq DEGs directly regulated by α1HCT, inferred by ATAC-seq, in hNPCs stably expressing α1HCT, with or without 20 mM KCl treatment.

(F, G, and H) Quantification by qRT-PCR of the DEGs' mRNA level in hNPCs stably expressing α1CCT, α1ACT, and α1HCT.

**Supplementary table 1. Voltage-gated calcium channel nomenclature, predicted C-terminal protein size, and stop codon locations**

| Protein Name | Alpha Subunit | Gene Name (human) | C-terminal Protein (Predicted Size) | Stop codon(s) location Amino Acid (Nucleotide) | Accession Number |
|---|---|---|---|---|---|
| Ca$_V$1.1 | α1S | *CACNA1S* | α1SCT (37 kDa) | Q141* (C508T); Q464* (C1477T) | NM_000069.3 |
| Ca$_V$1.2 | α1C | *CACNA1C* | α1CCT (70 kDa) | G230* (G1264T); I1011* (ATC3607-3609TAG) | NM_199460.3 |
| Ca$_V$1.3 | α1D | *CACNA1D* | α1DCT (60 kDa) | C333* (T1555A) | NM_001128839.3 |
| Ca$_V$1.4 | α1F | *CACNA1F* | α1FCT (50 and 60 kDa) | Q149* (C416T) | NM_005183.4 |
| Ca$_V$2.1 | α1A | *CACNA1A* | α1ACT (75 kDa) | P1846* (CCC5791-5793TAG) | NM_023035.3 |
| Ca$_V$2.2 | α1B | *CACNA1B* | α1BCT (60 kDa) | E93* (G429T) | NM_000718.4 |
| Ca$_V$2.3 | α1E | *CACNA1E* | α1ECT (55 kDa) | Q271* (C1039T) | NM_000721.4 |
| Ca$_V$3.1 | α1G | *CACNA1G* | α1GCT (55 kDa) | Q701* (C2846T) | NM_198382.3 |
| Ca$_V$3.2 | α1H | *CACNA1H* | α1HCT (70 kDa) | R1231* (C3939T, C3941A); S1234* (A3948T, C3950A) | NM_021098.2 |
| Ca$_V$3.3 | α1I | *CACNA1I* | α1ICT (75 and 85 kDa) | Q170* (C508T); Q440* (C1318T) | NM_001003406.2 |

**Supplementary table 2. Average percentage of subcellular localization**

| | Nuclear | | | Cytoplasm | | |
|---|---|---|---|---|---|---|
| Protein expression | Mean | SEM | N (# of fields of view) | Mean | SEM | N (# of fields of view) |
| α1S | 10.33 | 3.67 | 5 | 89.67 | 3.67 | 5 |
| α1SSTOP | 11.39 | 4.85 | 5 | 88.61 | 4.85 | 5 |
| α1C | 13.66 | 4.13 | 4 | 86.34 | 4.13 | 4 |
| α1CSTOP | 69.75 | 16.35 | 3 | 30.25 | 16.35 | 3 |
| α1D | 33.06 | 7.47 | 9 | 66.94 | 7.47 | 9 |
| α1DSTOP | 92.58 | 4.15 | 11 | 7.42 | 4.15 | 11 |
| α1F | 0.00 | 0.00 | 5 | 100.00 | 0.00 | 5 |
| α1FSTOP | 37.21 | 10.18 | 9 | 62.79 | 10.18 | 9 |
| α1A | 63.43 | 6.30 | 4 | 36.57 | 6.30 | 4 |
| α1ASTOP | 100.00 | 0.00 | 6 | 0.00 | 0.00 | 6 |
| α1B | 4.88 | 2.27 | 6 | 95.12 | 2.27 | 6 |
| α1BSTOP | 43.14 | 6.86 | 3 | 56.86 | 6.86 | 3 |
| α1E | 14.44 | 5.44 | 5 | 85.56 | 5.44 | 5 |
| α1ESTOP | 62.80 | 7.80 | 2 | 37.21 | 7.80 | 2 |
| α1G | 0.00 | 0.00 | 7 | 100.00 | 0.00 | 7 |
| α1GSTOP | 2.50 | 2.50 | 8 | 97.14 | 2.86 | 7 |
| α1H | 3.43 | 2.15 | 5 | 96.57 | 2.15 | 5 |
| α1HSTOP | 37.29 | 7.07 | 8 | 62.71 | 7.07 | 8 |
| α1I | 7.47 | 2.66 | 6 | 92.53 | 2.66 | 6 |
| α1ISTOP | 22.92 | 6.52 | 5 | 77.08 | 6.52 | 5 |

**Supplementary table 3. List of antagonists**

| Treatment | Concentration | Target | Type |
|---|---|---|---|
| AP5 | 100 μM | NMDA Channel | Antagonist |
| Nifedipine (Nif) | 10 μM | L-Type VGCC | Antagonist |
| W-7 Hydrochloride | 100 μM | Calmodulin | Antagonist |
| w-Agatoxin | 500nM | $Ca_v2.1$ | Antagonist |
| TTA-A2 | 100 μM | T-Type VGCC | Antagonist |
| 2-APB | 50 μM | IP3 Receptors | Antagonist |
| AP5 | 100 μM | NMDA Receptors | Antagonist |

**Supplementary table 4. α1CCT and α1ACT nuclear signal changes with different treatments**

| Treatment | Dose | Target | EV | α1CCT | α1CCT Percent Change | α1ACT | α1ACT Percent Change |
|---|---|---|---|---|---|---|---|
| Control | -- | -- | -3.1±0.6% | -0.02±0.04% | -- | -0.01±0.03% | -- |
| Glutamate (Glu.) | 100 μM | -- | -2.6±0.04% | +18.34±1.5% $p<0.0001$* | -- | -10.73±1.6% $p<0.0001$* | -- |
| AP5 | 100 μM | NMDA Channel | +1.0±0.05% | +11.39±0.014% | -6.95±0.014% $p<0.0001$ | -4.67±0.038% | +6.06±0.038% $p<0.0001$ |
| Nifedipine (Nif.) | 10 μM | L-Type VGCC | +1.7±0.04% | +3.1±0.014% | -15.24±0.014% $p<0.0001$ | -10.03 ± 0.020% | +0.7± 0.020% $p=0.7211$ |
| W-7 Hydrochloride | 100 μM | Calmodulin | +0.0+0.04% | +7.25±0.5% | -11.09±0.5% $p<0.0001$ | -5.76±0.9% | +4.97±0.9% $p=0.0321$ |
| ω-Agatoxin | 500 nM | $Ca_v2.1$ | +3.4±0.2% | +11.2±0.021% | -7.14±0.021% $p=0.0012$ | -12.67±0.021% | -1.94±0.021% $p=0.3551$ |
| TTA-A2 | 100 μM | T-Type VGCC | +2.2±0.6% | +12.81±0.020% | -5.53±0.020% $p=0.0071$ | -12.14±0.020% | -1.41±0.020% $p=0.4910$ |
| 2-APB and Ryanodine | 50 μM 100 μM | IP3 Receptors Ryanodine Receptors | -2.0±0.2% | +16.53±2.8% | -1.81±2.8% $p=0.5720$ | -13.49±0.018% | -2.76±0.018% $p=0.1293$ |

* Compared to control
Highlighted columns reflect the data reported in manuscript
